# Developmentally Delayed Epigenetic Reprogramming Underlying the Pathogenesis of Preeclampsia

**DOI:** 10.1101/2020.05.08.085290

**Authors:** Wei He, Yuan Wei, Xiaoli Gong, Luyuan Chang, Wan Jin, Ke Liu, Xinghuan Wang, Yu Xiao, Wenjing Zhang, Qiong Chen, Kai Wu, Lili Liang, Jia Liu, Yawen Chen, Huanhuan Guo, Wenhao Chen, Jiexia Yang, Yiming Qi, Wei Dong, Meng Fu, Xiaojuan Li, Jiusi Liu, Yi Zhang, Aihua Yin

## Abstract

Preeclampsia, a life-threatening pregnancy complication characterized by hypertension and multiorgan damage, affects 2-5% of pregnancies and causes 76,000 deaths per year. Most preeclampsia associated syndromes immediately dispel after removal of placenta, indicating a casual role of placenta in the pathogenesis. Failed transformation of spiral artery due to insufficient invasion and excessive apoptosis of trophoblast suggested developmental defects in preeclampsia placenta. However, the underlying molecular mechanisms that affected placenta development in preeclampsia remained elusive. Here we show that, in preeclampsia placenta, the epigenetic landscape formed during extraembryonic tissue differentiation was disrupted: dramatic chromatin accessibility shift affected known and novel genes implicated in preeclampsia. DNA methylation defects in preeclampsia affected lineage-defining PcG-controlled loci in trophectoderm. LTR12 retrotransposons associated with VCT/SCT-specific genes were hypermethylated. Meanwhile, hundreds of PcG-regulated EVT-specific gene promoters, which otherwise undergone post-ZGA extraembryonic-tissue-specific *de novo* methylation, were hypomethylated and hyper-activated. Together, these epigenetic defects resulted in placenta developmental delay in preeclampsia. The defective methylation pattern could be detected in serum cfDNA, and could be used to accurately predict preeclampsia at early pregnancy weeks in independent validation cohorts. Our data suggests that the preeclampsia placenta represents a stalled state of epigenetic reprogramming *en route* of development from trophectoderm to normal placenta.

---

Dramatic epigenetic transformation occurs during human embryogenesis^1–8^. After zygote formation, the overall epigenome was turned into an open, transcriptionally permissive environment^2^ by genome-wide removal of DNA methylation and H3K27me3 repressive mark^1,3,7^, until zygote genome activation (ZGA) when repressive epigenetic modifications were re-established on linage-specification genes by maternal and ZGA-active^6,9,10^ transcription factor priming, polycomb group protein (PcG) binding^11–16^, and *de novo* DNA methylation on CpG-island promoters^17^. In extraembryonic tissue (ExE), *de novo* DNA methylation on promoters of PcG-regulated genes primes trophoblast cell lineage for implantation. With defective development of placenta, insufficient invasion of extravillous trophoblast (EVT) to the maternal uterus decidua^18^ and excessive apoptosis of villous cytotrophoblast (VCT)^19^ could result in failed transformation of spiral arteries. The resulted insufficient supply of blood and hypoxia leads to reflective secretion of vasodilative factors such as s-FLT1 from placenta^20^, causing an avalanche of pathological events leading to hypertension and multiorgan failure in preeclampsia^21^. To understand how placenta development is disrupted in preeclampsia, we assessed single cell and bulk genome-wide DNA methylation, histone modification and chromatin accessibility from different stages of human embryonic development to placenta, including single gamete (sperm and oocyte), zygote, 2-cell stage, 4-cell stage, 8-cell stage, morula, inner cell mass and trophectoderm from blastocyst stage, primed embryonic stem cells (ESC), trophoblast cells and preeclampsia- or non-preeclampsia placenta using a collection of data^1–4,6,22–24^ (Extended Data Table S1a and S4).

## Chromatin accessibility and TFBS changes in preeclampsia placenta

Global chromatin accessibility landscapes from placenta of preeclampsia pregnancy is drastically different from normal ones (Extended Data Figure 1). Statistically significant difference on chromatin accessibility was identified on 3,834 genomic loci, in which 1,960 peaks were gained in preeclampsia (preeclampsia-gain) and 1,874 were lost in preeclampsia (preeclampsia-loss) (Extended Data Table S2). These preeclampsia-specific, differentially regulated loci were particularly enriched for active TSS, CpG-islands, polycomb regulated regions and enhancers^11,12^ (Figure 1a), suggesting a possibility that these regions were under control of DNA methylation ^25^. Particularly, more activating regions (TSS, Enhancer) were repressed, and more repressive regions (polycomb controlled heterochromatin and repressed regions) were found open in preeclampsia (Figure 1a). Peaks on preeclampsia-gain loci were usually found in pre-ZGA to early-ZGA embryonic stages, and preeclampsia-loss loci were more frequently found in ZGA to post-ZGA (Figure 1b). Preeclampsia-loss loci associates with essential transcription factors such as *PAX6, GATA3*, as well as the imprinted genes *H19*^26^ and *PEG3*^27–29^, suggesting defective DNA methylation, genetic imprinting and transcriptional activation (Figure 1c). Many known preeclampsia-associated genes such as *AGT*^30–32^ (Figure 1e,1f), *FLT1* ^33–38^, *PAPPA2*^39,40^, *KDR*^41^, and *KISS1R*^42^ were found to be associated with significantly upregulated chromatin accessibility (Figure 1d). In concordance to previous studies ^36,37,20,39^, chromatin accessibility changes occurred in cis-regulatory regions for genes (Extended Data Figure 2) implicated in preeclampsia etiology as biomarkers^20,39,43^, conferred genetic susceptibility^33–35^, or with direct functional link^38,40,41,44,45^. Interestingly, some of these genes were mis-activated by activating enhancers with otherwise ‘poised’ bivalent chromatin with both repressive and activating histone marks, as shown in the case of *AGT* (Figure 1f), suggesting a possible role of PcG in preeclampsia.

**Figure 1:**
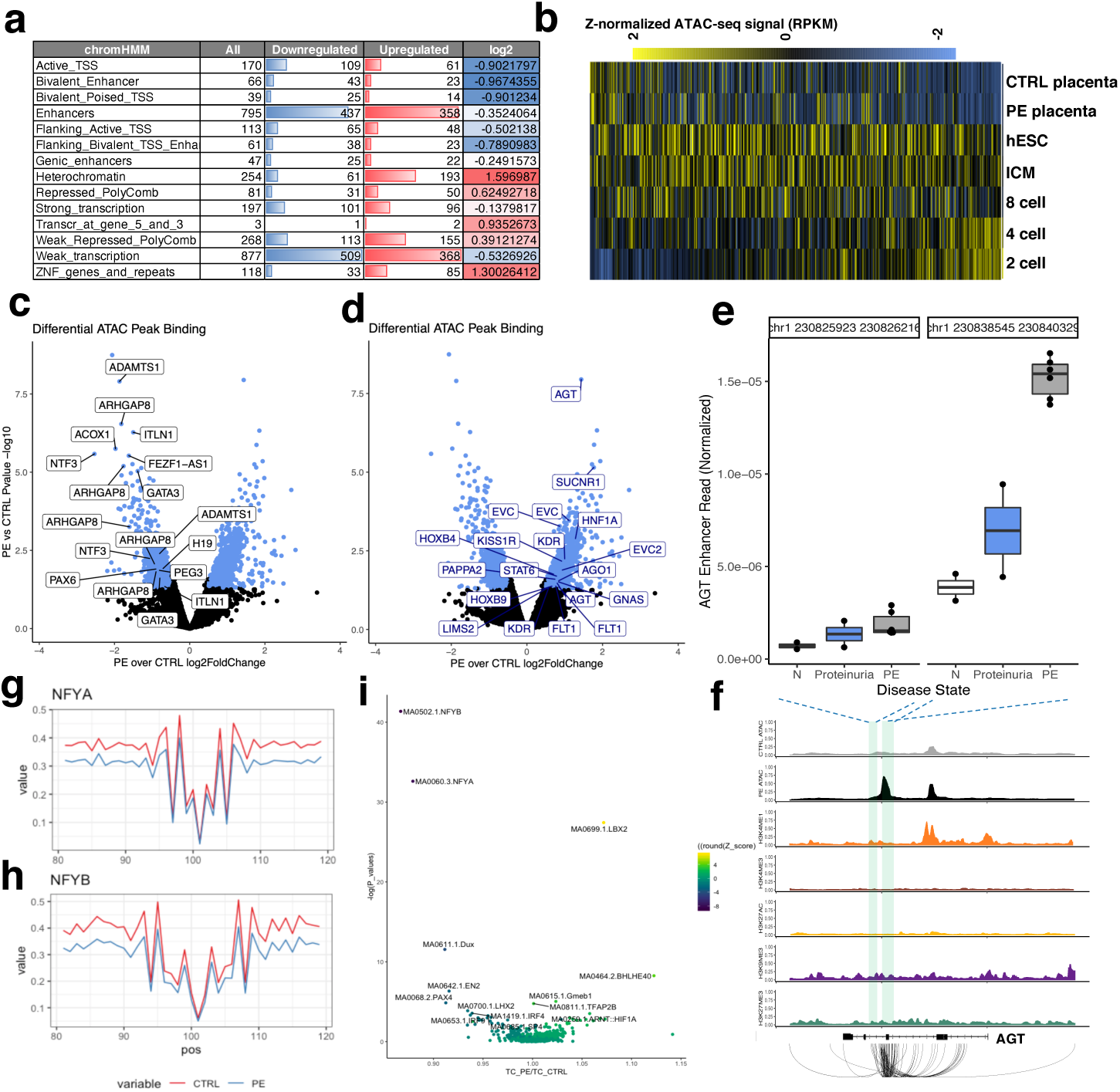
Preeclampsia-specific placental chromatin accessibility defects. a: Enrichment of significantly different ATAC-seq peaks on different classes of^11,12^ chromHMM genomic regions. b: Z-normalized ATAC signal (RPKM) intensity across embryonic stages on IDR peaks showed differences between preeclampsia and non-preeclampsia placenta. c: Differential ATAC binding (x: PE/CTRL log2 fold change; y: logP value (base10)) with names of significantly down-regulated ATAC peaks associated genes. d: Differential ATAC binding (x: PE/CTRL log2 fold change; y: logP value (base10)) with names of significantly up-regulated ATAC peaks associated genes. e: AGT enhancer chromatin accessibility is significantly increased in placenta of preeclampsia pregnancy or proteinuria (without GHT). y: normalized ATAC-peak coverage. f: Raw ATAC-seq track of normal (CTRL-ATAC) and preeclampsia (PE-ATAC) placenta, as well as ChIP-seq track of placenta (Roadmap) and chromatin interaction (4DGenome) showed that a strong cryptic AGT enhancer which is usually *not* activated in placenta is strongly activated in preeclampsia placenta. g: Transcription factor footprint of NFYA is significantly decreased in preeclampsia. X axis: position (100=TF binding site) around the TF binding site for NFYA; Y axis: normalized read depth; Red: non-preeclampsia (CTRL) placenta; Blue: preeclampsia (PE) placenta. h: Transcription factor footprint of NFYB is significantly decreased in preeclampsia; X axis: position (100=TF binding site) around the TF binding site for NFYB; Y axis: normalized read depth; Red: non-preeclampsia (CTRL) placenta; Blue: preeclampsia (PE) placenta. i: Scatter plot denoting the ratio of TF footprint strength for preeclampsia over normal placenta (X-axis) and P-value (Y-axis) for TF binding differences on placenta. X axis: Difference of read depth ratio of preeclampsia-vs.-non-preeclampsia placenta; Y axis: log P value. Color hue: Z from -8 to 4.

Functional enrichment of preeclampsia-gain and -loss regions suggested significant association of VEGFR, FGFR, TGF, WNT and MET signalling pathway involvement, indicating that the promiscuous oncogenic pathways activating *de novo* methylation during extraembryonic tissue development^17^ might be affected (Extended Data Figure 2). Furthermore, cell death related pathways such as lipophagy and apoptosis were also associated with these regions. Finally, transcription factor footprinting analysis^46,47^ shows that binding of NFYA and NFYB, two key transcription factors might regulate zygotic genomic activation (ZGA)^6,9^, were down-regulated in preeclampsia placenta (Figure 1g-i). Binding of another known ZGA-associated transcription factor DUX4 was also found to be decreased in some but not all of the cases (Figure 1i). Overall, these data suggested that preeclampsia placenta was associated with a defect in epigenetic reprogramming.

## Methylation changes in early onset preeclampsia placenta on polycomb binding site in trophectoderm

As suggested by ATAC-seq data, DNA methylation related changes were prevalent in preeclampsia placenta. We went on to analyze genome-wide methylation from fetal and maternal surface of placentas. To this end, we identified 4,418 differentially methylated regions (DMR), in which 2,710 were hypomethylated (preeclampsia-hypo) and 1,271 were hypermethylated (preeclampsia-hyper) in preeclampsia placenta (Extended Data Table S3 and Extended Data Figure 3). DNA reads on DMR were classified into high-methylation (methylated) and low-methylation (demethylated) haplotype based on the overall methylation frequency they carried (Extended Data Figure 4). On most DMR, the difference between preeclampsia and normal placentas was dominated by one single class of methylation haplotype.

Mean methylation level on these DMR distinguished all early-onset severe preeclampsia (EOSPE) placenta and a majority of late-onset preeclampsia (LOPE) placenta from other non-preeclampsia placenta from normal donors or donors of other pregnancy complications, suggesting that preeclampsia is a biologically distinct disease (Figure 2a and Extended Data Figure 5). Hierarchical clustering of the DMR showed clusters of preeclampsia-hyper and preeclampsia-hypo DMR behaving consistently across samples, suggesting these DMR are under control of similar molecular mechanisms (Figure 2a).

**Figure 2:**
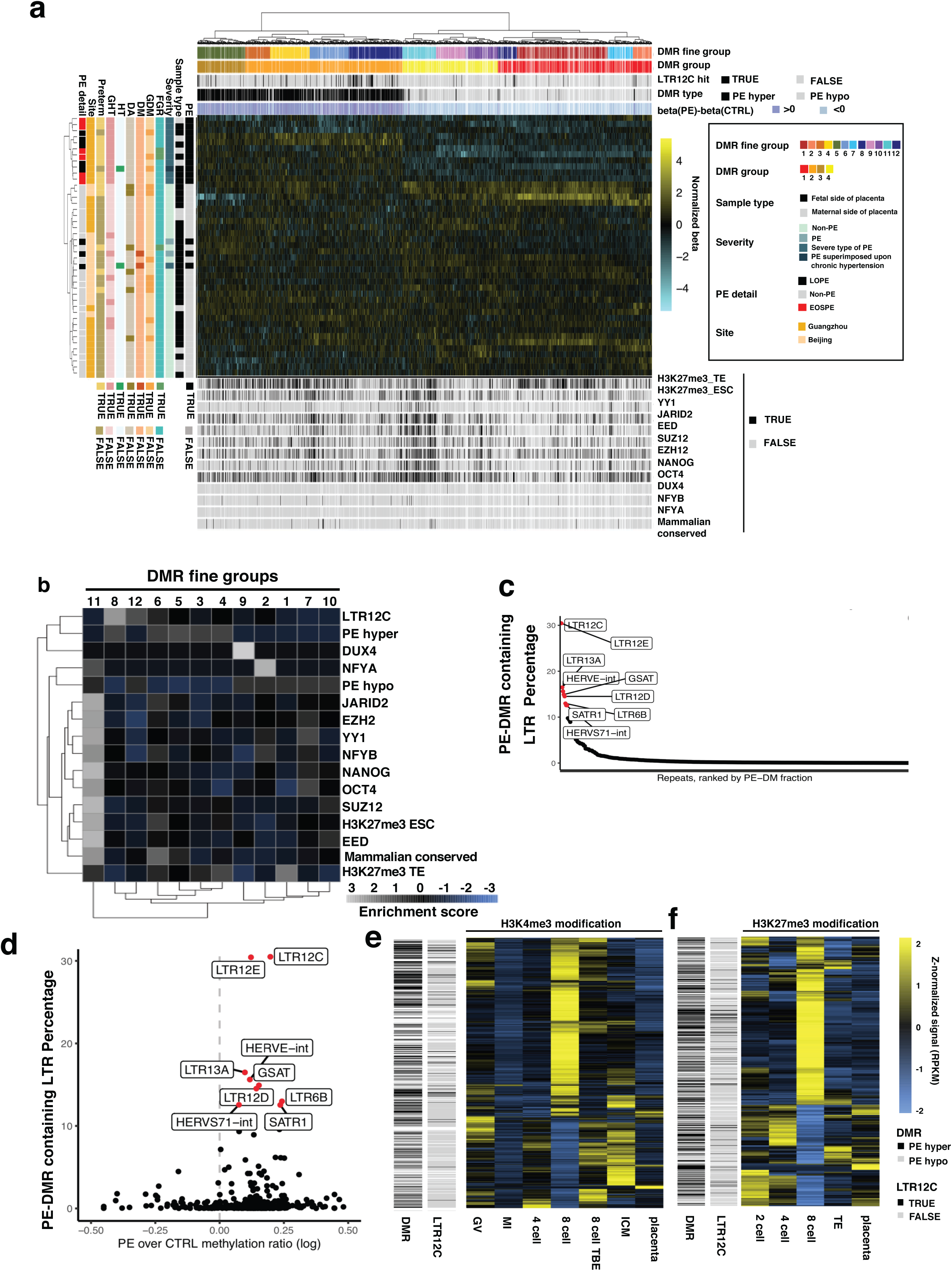
Bimodal genome-wide methylation defects associated with preeclampsia. a: Heatmap of normalized beta value of preeclampsia DMR including both preeclampsia-hyper (PE-hyper) DMR and preeclampsia-hypo (PE hypo) DMR across 43 placenta samples from preeclampsia (PE), fetal growth restriction (FGR), gestational diabetes mellitus (GDM), gestational hypertension (GHT), chronic hypertension (HT), chronic diabetes mellitus (DM), preterm, diamniotic pregnancy (DA) pregnancies, or pregnancies without above complications. Preeclampsia placentas can be distinguished from non-preeclampsia placentas based on DNA methylation profile. Unsupervised hierarchical clustering showed clear divergence of PE-hyper and PE-hypo DMR across samples. PE-hyper DMR were enriched with LTR12C elements. A subgroup of PE-hypo DMR was highly enriched with PRC2 binding (EZH2/EED/JARID2/SUZ12/YY1), H3K27me3 modification in ESC and trophectoderm, and pluripotency factor (OCT4/NANOG) binding. b: Enrichment of transcription factor binding, H3K27me3 modification in trophectoderm (TE) and ESC, LTR12C, and methylation differences (PE-hyper and PE-hypo) in DMR suggesting that group 6 and group 11 DMR were regulated by PRC2. Group 11 DMR is also associated with NFYB/NFYA, suggesting a functional link between transcription factor binding defect and methylation defect. Group 9 is associated with DUX4 binding, and Group 2 is associated with NFYA binding. Group 8/12/6 were associated with LTR12C. c: LTR12 retrotransposon family were the most affected repeat element in preeclampsia placenta. Y-axis: Percentage of repeat element containing preeclampsia-specific DMR; X-axis: rank of percentage from high to low; Red dots with names are repeats with significant enrichment (Fisher’s exact test P<0.05) of DMR d: LTR12 family retrotransposon are hypermethylated in preeclampsia placenta. X-axis: log-ratio of mean methylation level of preeclampsia placenta over normal placenta for individual class of repeat element; Y-axis: Fractions of repeats containing preeclampsia-specific DMR. Red dots with names are repeats with significant enrichment (Fisher’s exact test P<0.05) of DMR. e: Z-normalized H3K4me3 Cut-and-Run signal (RPKM) within +-100bp of preeclampsia-specific DMR across different embryonic stages and placenta samples showed transcriptional-dependent enrichment of H3K4me3 mark on preeclampsia-hyper and LTR12C DMR in 8-cell-stage. PLC: Placenta; ICM: inner cell mass; TBE: transcriptional block; MI: meiosis I oocyte; GV: geminal vesicle. f: Z-normalized H3K27me3 Cut-and-Run signal (RPKM) +-10000bp of preeclampsia-specific DMR across different embryonic stages and placenta samples showed exclusion of H3K27me3 mark on preeclampsia-hyper and LTR12C DMR in trophectoderm. PLC: Placenta; TE: trophectoderm. Black lines: LTR12C-contained DMR; Red lines: preeclampsia-hyper DMR;

Transcription factor binding sites were enriched in groups of preeclampsia-hypo DMR (Extended Data Figure 6). Correlation of known TFBS on the DMR showed that NFYA and NFYB bind to a group of preeclampsia-hypo DMR (group 11) (Figure 2b), which are also enriched with known ESC binding sites of PRC1 (CBX1/CBX2/CBX3), PRC2 (EED/EZH2/JARID2/SUZ12), PRC recruiter YY1, and the master regulator of pluripotency (OCT4/NANOG), whilst overlapping significantly with H3K27me3 in both ESC and trophectoderm. Gene pathway enrichment analysis showed that group 11 DMR are significantly associated with lineage specification genes (Extended Data Figure 7), suggesting that this group of DMR are likely to be regulatory regions under consistent control of PcG during extraembryonic tissue development. As expected, these regions were also enriched for evolutionarily conserved ExE-*de-novo* methylated region (Figure 2a and 2b).

PcG is part of the molecular machinery required for maintaining stable cell fate in differentiated lineages during embryogenesis. The genetic evidences for its role in memorizing cell fate initially came from the *Drosophila* polycomb mutants, whose developmental defects were phenotypically apparent only after body plan formation^48–52^. Biochemically, PcG maintains lineage-specification gene silencing^53–56^ and gene imprinting^28,57–59^ by binding to loci with PRE motif containing demethylated CpG DNA and decorated by H3K27me3 marks, and establishing DNA methylation on neighboring CpG shores^60,61^. In placenta, genetic defects affecting PcG activities, particularly YY1 and its imprinted binding partner SFMBT2^59^, result in highly specific developmental defects such as hydatidiform moles^62^, placenta growth restriction^63^ or enlargement^64^. The fact that a number of lineage-specification gene-associated, PcG-bound loci were hypomethylated in preeclampsia but not normal placenta suggests that cell lineage specification might be affected in preeclampsia. We also found that H3K27me3 in trophectoderm but not ESC were enriched on other groups (group 1 and group 4) of DMR, and group 6 DMR were marked with ESC H3K27me3 and bound with PRC to a lesser extent, suggesting that PRC might be recruited to these groups of DMR during trophectoderm differentiation. Both evolutionarily conserved (Extended Data Figure 8a-8b) or human-specific (Extended Data Figure 8c-8d) preeclampsia-hypo regions were highly methylated in both normal placenta and human cancer (Extended Data Figure 8a, 8c) but not in preeclampsia placenta (Extended Data Figure 8b, 8d), suggesting that the innate developmental trajectory of extraembryonic tissue, which is mirrored by cancer, is defective in preeclampsia.

On the contrary, preeclampsia-hyper DMR does not distinguish cancer to normal samples (Extended Data Figure 8e and 8f). Enrichment analysis showed that whilst preeclampsia-hypo DMR were enriched with UTR and cis-regulatory elements (Extended Data 9a, 9b) the preeclampsia-hyper DMR were enriched with retrotransposons with long terminal repeat (LTR) (Extended Data Table S3 and Extended Data Figure 9). Among retrotransposons, the primate-specific LTR12 family^23^ were dominated by preeclampsia-hyper DMR (Figure 2a-2d). Up to 30% of LTR12C and LTR12D retrotransposons contain preeclampsia-hyper DMR (Figure 2c), and most LTR12C-contained DMR are hypermethylated in preeclampsia placenta (Figure 2d).

LTR12 was known to be hypomethylated in primate sperm^23^ and was considered as a primate-specific innovation of imprinting. We found that LTR12 hypomethylated in human sperm were commonly hypermethylated in preeclampsia (Extended Data Figure 9c-9e). The methylation level of LTR12 family distinguishes preeclampsia and normal placenta with high specificity (Extended Data Figure 10).

To further understand the underlying molecular mechanism of differential methylation in preeclampsia, we analyzed histone modification profiles on these preeclampsia-associated DMR using existing Cut-And-Run histone modification data from different embryonic stages^2,4^. H3K4me3 modification on preeclampsia-associated DMR showed bimodal changes at early-ZGA stage (Figure 2e), with more H3K4me3-positive preeclampsia-hypermethylated regions (including LTR12C) and H3K4me3-negative preeclampsia-hypomethylated regions. The landscape of H3K4me3 on preeclampsia-hypermethylated regions were established with active transcription, as the evolution of H3K4me3 modification pattern was from 4-cell stage to 8-cell stage in transcriptionally blocked embryo (TBE) ^2^ (Figure 2e). For H3K27me3, transient enrichment was also found in 8-cell stage (Figure 2f). Finally, differential chromatin accessibility were detected on DMR. Such loci were enriched with known pathways for placenta development and function such as NOTCH3, VEGFA, MET, FOXO, ECM proteoglycan regulation, and coagulation (Extended Data Figure 19). Together, these results suggest that DNA methylation defects in preeclampsia occurs on lineage-specifying genes under active regulation by PcG during trophectoderm-placenta development.

## Preeclampsia-hypermethylated regions are associated with VCT/SCT-specific genes and ZGA-active LTR12 retrotransposon

Fetal extraembryonic tissue builds placenta by providing villous cytotrophoblast (VCT)^65^ which forms the major structure of fetal face of placenta, and differentiates along divergent trajectories towards either syncytiotrophoblast (SCT)^66^, or extravillous trophoblast (EVT)^66^. Successful transformation of spiral artery requires coordinated, efficient generation of trophoblasts of both lineages. Using trophoblast single cell RNA sequencing data^22^, we built a pseudotime differentiation trajectory of VCT towards EVT or SCT using DMR-associated genes (Figure 3a and Extended Data Figure 11). Preeclampsia-hyper DMR associated genes were highly expressed in VCT (Figure 3a, large) and to a lesser extent SCT and decidua EVT, but not in the placental EVT, suggesting preeclampsia-hyper DMR might regulate VCT-specific or SCT-related genes. Concordant to this finding, on the fetal side of placenta, we found that the methylation haplotype frequency and overall methylation level of preeclampsia-hyper LTR12C DMR, could differentiate fetal side but not maternal side of placentas of preeclampsia pregnancy from normal ones (Figure 3b), suggesting the cell type enriched in fetal side of placenta as the major contributor of the methylated haplotypes in preeclampsia-hyper DMR. Together, these results suggest preeclampsia-hyper DMR might implicate in VCT (and SCT) function.

**Figure 3:**
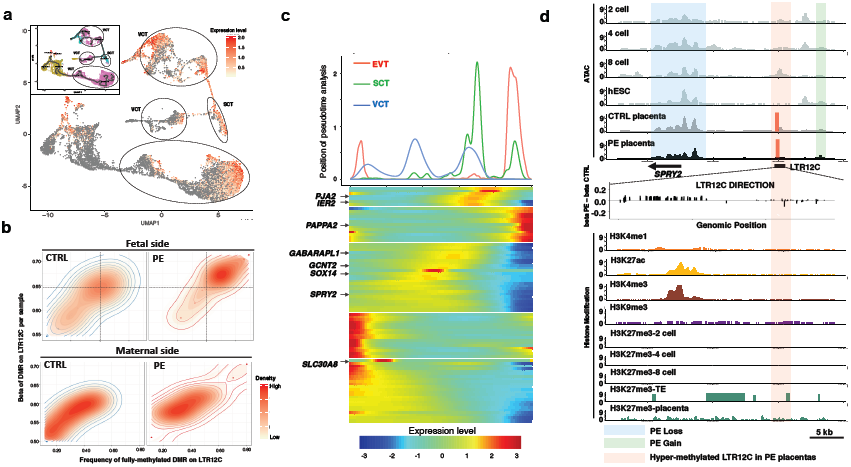
Genomic region hypermethylated in preeclampsia placenta are associated with VCT- and SCT-specific genes and ZGA-active retrotransposon. a: UMAP projection and pseudotime trajectory (black) of single-cell RNA sequencing results of trophoblasts. Heat color denotes normalized sum of preeclampsia-hyper DMR associated gene expression level in each cell. Inset: color-coded classes of trophoblasts. pink: VCT; blue: SCT; yellow: EVT; red: decidua EVT. b: Preeclampsia-hyper DMR methylation level (Y) and fully-methylated DNA read frequency (X) are different between preeclampsia and normal placenta on the fetal, but not maternal surface. Individual placenta samples were denoted as dots, and 2-D distribution were outlined. c: Gene expression level modelled on pseudotime trajectory for preeclampsia-hyper DMR associated genes. Top: distribution of cells at different timepoints (x-axis) of pseudotime. Blue: VCT; Red: EVT; Green: SCT. d: ATAC-seq of 2-cell, 4-cell, 8-cell, primed-ESC, preeclampsia (PE) and normal (CTRL) placenta on *SPRY2* loci (upper panel), associated with H3K9me3 (purple), H3K27me3 (green), H3K4me1 (orange), H3K27ac (yellow) and H3K4me3 (brown) modification ChIP-seq signal (lower panel). Red lines denote DMR location. Pink shadow denotes preeclampsia-hypermethylated LTR12C in *SPRY2* which is transcriptionally activated at early-ZGA (8-cell). Green shadow denotes ATAC peaks gained in preeclampsia placenta (PE gain). Blue shadow denotes ATAC peaks lost in preeclampsia placenta (PE loss). Middle panel: methylation level difference of CpG in the LTR12C element, between preeclampsia and normal placenta. Direction of LTR12C were denoted as arrow.

We used Monocle^67^ to infer preeclampsia-hyper DMR associated gene expression level along the trophoblast differentiation trajectories (Figure 3c). By projecting cells on this pseudotime evolution trajectory, we found a subset of genes with associated LTR12C containing preeclampsia-hyper DMR were selectively expressed in VCT. During embryonic development, transcription from these LTR12C were selectively activated at ZGA (8-cell stage) (Figure 1b, 3d and Extended Data Figure 12), when the NFYA/NFYB transcription factors^6,9^ were most active.

Methylation differences between preeclampsia and normal placentas on these LTR12C elements were very similar (Extended Data Figure 12), with hypermethylation at the 5’ end of retrotransposon. These sites are demethylated in pre-ZGA stages and re-methylated post-ZGA (Extended Data Figure 13). Hence, hypermethylation of these sites suggested a possibility that these retrotransposons were dysregulated before or during ZGA.

Many of the LTR12C hypermethylated in preeclampsia were with permissive histone mark in early ZGA and repressive histone mark re-established in trophectoderm (Figure 2e and 2f). For example, in *SPRY2* (Figure 3d), *PAPPA2, CDKN3* and *YY1* (Extended Data Figure 14), the hypermethylated LTR12C was active during 8-cell stage and repressed in trophectoderm. Analyzing chromatin interaction between the cognate promoters of these genes and the hypermethylated LTR12C showed an anti-correlation between LTR12C hypermethylation and transcriptional activation: *PAPPA2*, which is upregulated in preeclampsia^39^ (Extended Data Figure 14a), was associated with a hypermethylated LTR12C in its adjacent topological association domain (TAD) as suggested by chromatin interaction frequency; in contrast, *SPRY2, CDKN3*^68^ and *YY1*^69^ which are downregulated in preeclampsia, were associated with a hypermethylated LTR12C in their own TAD. These results suggest that preeclampsia-specific hypermethylation of LTR12C might result in down-regulation of transcriptional activity in its own TAD, and might up-regulate transcriptional activity in the adjacent TADs.

## Post-ZGA-active, embryonically protected, PcG-controlled CpG island promoters associated with EVT-specific genes were hypomethylated

We used a similar strategy to locate the cell type(s) most affected by preeclampsia-hypo DMR. Preeclampsia-hypo DMR associated genes were exclusively expressed in EVT (Figure 4a), without significant expression in either SCT or VCT. Meanwhile, methylation levels of preeclampsia-hypo DMR could differentiate the maternal side, but not fetal side, of preeclampsia placenta from normal ones (Figure 4b). Pseudotime trajectory mapping also showed preeclampsia-hypo DMR associated gene expression were elevated along the evolution towards EVT (Figure 4c). The preeclampsia-hypo DMR associated with these genes are generally located in the promoter or enhancer elements, and are closely associated with regions with preeclampsia-gain of chromatin accessibility (Extended Data Figure 15).

**Figure 4:**
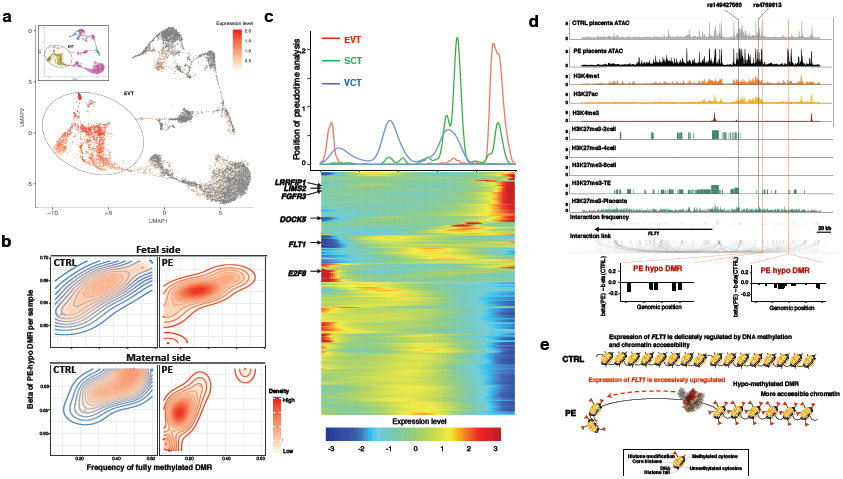
Post-ZGA-active, embryonically protected CpG island promoters associated with EVT-specific genes were hypomethylated in preeclampsia placenta. a: UMAP projection and pseudotime trajectory (black) of single-cell RNA sequencing results of trophoblasts. Heat color denotes sum of preeclampsia-hypo DMR associated gene expression level in each cell. Inset: color-coded classes of trophoblasts. pink: VCT; blue: SCT; yellow: EVT; red: decidua EVT. b: Preeclampsia-hypo DMR methylation level (Y) and fully-methylated DNA read frequency (X) are different between preeclampsia and normal placenta on the maternal, but not fetal surface. Individual placenta samples were denoted as dots, and 2-D distribution were outlined. c: Gene expression level modelled on pseudotime trajectory for preeclampsia-hypo DMR associated genes. Top: distribution of cells at different timepoints (x-axis) of pseudotime. Blue: VCT; Red: EVT; Green: SCT. d: ATAC-seq of preeclampsia (PE) and normal (CTRL) placenta, associated with H3K4me3 (brown), H3K27ac (yellow), H3K4me1 (orange) modification ChIP-seq signal on *FLT1* loci. H3K27me3 modification (green) (Cut-and-Run for embryo, and ChIP-seq for placenta) at different developmental stages on *FLT1* loci shows preeclampsia-gained enhancer was under control of PcG and marked by H3K27me3 at trophectoderm stage. Red dashed line denotes preeclampsia-hypo DMR location. Positions for variants associated with increased risk for preeclampsia were denoted. Middle panel: Chromatin interaction intensity (gray histogram) and individual chromatin interaction links (bottom curves) on the region, showing that the preeclampsia-gained enhancer interacts with poised and regular promoter of *FLT1*. Lower panel: methylation difference of CpG in the preeclampsia-hypo DMR, between preeclampsia and normal placenta. e: Schematic drawing for epigenetic mechanism controlling *FLT1* overexpression in preeclampsia.

Gains of chromatin accessibility of these genes were exclusively found in placenta, but not earlier developmental stages (Extended Data Figure 15), implicating that these chromatin openings formed no earlier than TE-to-ExE transition. Since ExE-specific *de novo* methylation of promoters is non-cell-autonomous ^17^, preeclampsia-specific hypomethylation on these genomic loci implies a failure of *de novo* methylation during TE-to-ExE transition, which in turn suggests an early defect at the time of implantation and failed oncogenic extracellular signalling pathway activation^17^.

In all EVT-expressed genes, the VEGF receptor *FLT1* is of particular interest because it was biochemically^20^ and genetically^33^ linked to preeclampsia, and was found to be directly causing a preeclampsia-like phenotype in transgenic animal model downstream of endometrium VEGF^38^. We identified preeclampsia-gained enhancers for *FLT1* with gain of chromatin accessibility and loss of DNA methylation on H3K27ac-marked loci (Figure 4d). These enhancer encompassed the known fetal genome susceptibility locus rs4769613 and rs12050029 for preeclampsia^33^ and were conformationally linked to alternative *FLT1* promoters (Figure 4d). Moreover, these enhancers were marked with re-established H3K27me3 in trophectoderm (Figure 4d), suggesting that *FLT1* were directly targeted by PcG^58^ during TE-to-ExE transition and loss of such regulation may result in s-FLT1 overexpression (Figure 4e).

## Stalled epigenetic reprogramming of placenta in early onset preeclampsia

Together, we conclude that specific epigenomic changes in preeclampsia placenta occured on linage-specification related chromatin loci, leads to dysregulation of ZGA-active LTR12 family retrotransposon, and post-ZGA defect during TE-to-ExE transformation. These observations lead us to hypothesize that whether failed epigenetic reprogramming underlied the defective development of placenta in preeclampsia^70^. We tested this hypothesis by projecting placenta bulk methylation or ATAC sequencing data onto the evolution landscape built with single cell sequencing data. Mapping DNA methylation data positioned preeclampsia placenta along the innate developmental trajectory from trophectoderm cells towards trophoblast, whilst normal placenta clustered with *ex vivo* induced trophoblast (Extended Data Figure 16). Similarly, using chromatin accessibility on DMR regions (Extended Data Figure 17), we found that unlike the normal placenta which clustered together with endogenous trophoblast, preeclampsia placenta was positioned along the trajectory from trophectoderm towards trophoblast (Figure 5). Altogether, these results suggest that preeclampsia placenta is developmentally delayed not only anatomically, but also epigenetically.

**Figure 5:**
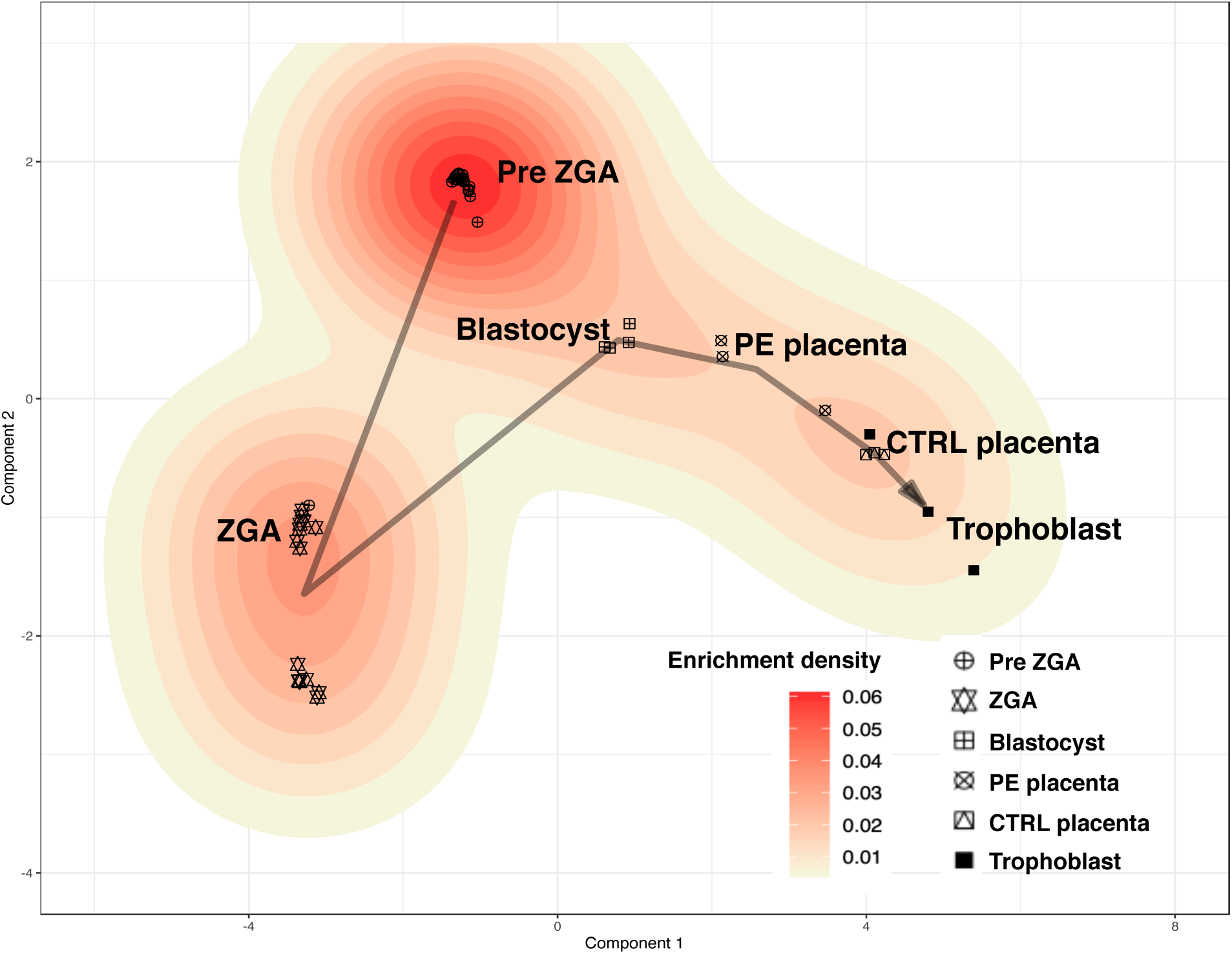
Stalled placenta epigenome development in preeclampsia. UMAP projection and pseudotime trajectory for single cell and bulk tissue ATAC sequencing from different embryonic stages to placenta, showing that preeclampsia placentas are stalled *en route* of the trajectory from blastocyst to trophoblast.

## *In vivo* evidence for the early stalled development of placenta from cell-free DNA

So far, we draw analysis based on sequencing data from term placenta. Due to continuous post-implantation development, these data might not accurately reflect the epigenomic landscape at earlier gestational stages. Serum cell-free DNA (cfDNA) originated from apoptotic cells in human body^71^. In early stages of pregnancy, cfDNA from pregnant female contains DNA of fetal origin^72^, which retains their original covalent chemical modification^73,74^. We hypothesized that sequencing cfDNA methylation profile from pregnant female, at very early gestational age, might be used to deduce the placental developmental status. If so, sequencing cfDNA methylation profile at early gestational weeks might also predict pregnancy outcome.

We sequenced the DNA methylation profiles on preeclampsia-associated DMR from cell-free DNA from 15-20w GA (gestational age) blood draws of 22 preeclampsia or normal pregnant females and 10 nongravida females using a capture panel (Extended Data Table S6). Analyzing the methylation haplotypes carried by sequencing reads showed that normal placenta associated (Figure 6a) methylation haplotype on these DMR could be readily detected in cfDNA of female of normal, but not preeclampsia, pregnancy (Figure 6a). Similarly, preeclampsia-placenta associated methylation haplotypes could be only detected in samples from pregnancy females who latter developed preeclampsia (Figure 6b and Extended Data Figure 18). These results suggest that DNA methylation defects observed in preeclampsia placenta occurred in early gestational week.

**Figure 6:**
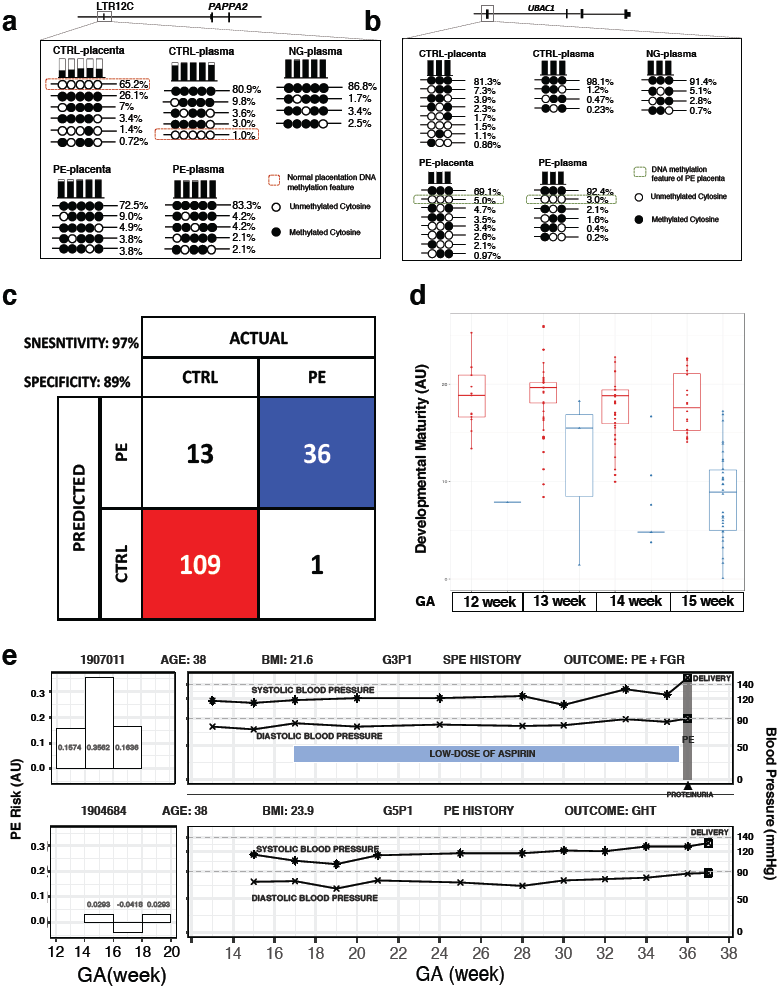
*In vivo* evidence for the early stalled development of placenta in cell-free DNA. a: Normal pregnancy associated methylation haplotype on a preeclampsia-hyper DMR on LTR12C element 5’ of *PAPPA2* could be detected in cell-free DNA. Mean methylation level (beta: top columns. black: methylated, white: unmethylated), individual CpG-loci methylation combinations on DNA sequencing reads (methylation haplotype, bottom panels) and associated frequency of each methylation haplotype (figures on the right) were shown for control placenta, preeclampsia placenta, control pregnant female, preeclampsia pregnant female, and nongravida female. Red dashed box denotes the normal pregnancy associated methylation haplotype which could only be found in placenta or cell-free DNA from control pregnant female. b: Preeclampsia pregnancy associated methylation haplotype on a preeclampsia-hypo DMR 5’ of *UBAC1* could be detected in cell-free DNA. Green dashed box denotes the preeclampsia pregnancy associated methylation haplotype which could only be found in placenta or cell-free DNA from preeclampsia pregnant female. c: Sensitivity and specificity in blinded validation for predicting preeclampsia with cell-free DNA methylation haplotype in a retrospective cohort. d: Post-hoc analysis showing predicted placenta developmental maturity is lower in preeclampsia individuals at early GA weeks. (Y-axis) Predicted placenta developmental maturity with cell-free DNA methylation haplotype from female who latter developed preeclampsia (blue) or normal (red). (X-axis) GA weeks. e. Prospective prediction of preeclampsia risk (Y-axis) from consecutive blood draws of two volunteers (left two panels) and clinical outcome of the two volunteers (right two panels), showing that the methylation haplotype deduced preeclampsia risk could predict final clinical outcome.

Because these methylation haplotypes were never observed in nongravida female, they represent placenta-specific biomarkers which could be used to non-invasively track placental development. To validate that early gestational week cfDNA methylation profile could be used to predict preeclampsia in later stage, we conducted a double-blinded, retrospective validation experiment. We firstly built a general linear model to convert the sequenced cfDNA methylation profile as a quantitative measure of placenta development to predict pregnancy outcome, which showed a 100% accuracy to predict preeclampsia in this small cohort. Blood draws collected at 13-20w GA, originally for NIPT, from 159 pregnant females, in which 37 were of preeclampsia, were used to validate the assay. Methylation profiles of these cfDNA were captured and sequenced.

Pregnancy outcome prediction based on methylation haplotype achieved a 97% sensitivity and 89% specificity in this cohort (Figure 6c). Post-hoc analysis showed that, starting from 13w GA, the cfDNA methylation profile from females who latter developed preeclampsia was significantly delayed compared to normal ones (Figure 6d).

Finally, we prospectively collected blood draws from two females at their early pregnancy to perform methylation profiling. Consecutive blood draws from 13-17w GA showed stable methylation profile measurement (Figure 6e, left panel), suggesting that the reprogramming of placental epigenome might be largely finished at 13w GA. Furthermore, we found the measurement accurately predicted their pregnancy outcome (Figure 6e, right panel).

## Discussion and Conclusion

Placenta, as the maternal-fetal interface, is essential for fetal development. Although it was known for decades that anatomical defective placental development underlies the pathogenesis of serious pregnancy complications such as fetal growth restriction and preeclampsia, their underlying molecular mechanism are largely unknown. Here we showed that preeclampsia is a disease associated with developmentally delayed placenta accompanied with failed epigenetic reprogramming characterized by specific DNA methylation and chromatin accessibility defects which together affected development of trophoblasts, and stalled the evolution of preeclampsia epigenome along its innate developmental trajectory. Our results not only provided insights into preeclampsia etiology, but also provided a mean for accurate identification of these high risk female at early GA weeks.

## Supporting information

Supplementary Data 1a Placenta Characteristics

Supplementary Data 1b Placenta Donor Characteristics

Supplementary Data 2 ATAC-seq differential peaks

Supplementary Data 3 Differentially methylated regions

Supplementary Data 4 Data used in the paper

Supplementary Data 5 Software used in the paper

Supplementary Data 6 Retrospective Clinical Cohort Characteristics

## Author Contributions

Y.Z, and A.H.Y conceived and designed the study. W.H., Y.W., X.L.G., J.X.Y, and Y.M.Q. collected clinical placenta and serum samples. Y.Z., K.L., X.H.W., and Y.X. arranged and collected clinical tumor and normal tissue samples. W.H. collected the clinical data for pregnant female, and set blinding for the validation cohort. W.J. collected the prospective prediction samples and their associated clinical data, performed table statistics on clinical data. Y.Z., Q.C., W.H.C., K.W. and H.H.G. developed the *Tequila 7N* single-stranded methylation sequencing assay. Y.W., X.L.G., W.J., M.F., W.D. and K.W. collected, dissected, and processed placenta samples for ATAC-seq. K.W., L.L.L., L.Y.C., Q.C., J.L., Y.W.C. and W.H.C. collected, dissected, and processed placenta samples for methylation capture sequencing. K.W., L.L.L., L.Y.C., Q.C., J.L., Y.W.C., H.H.G., Y.X. and W.H.C. collected, dissected, and processed tumor and normal tissue samples for methylation capture sequencing. K.W., L.L.L., L.Y.C., Q.C., J.L., Y.W.C. and W.H.C. processed plasma samples for methylation capture sequencing. Y.Z., L.Y.C., W.J.Z. and X.J.L. designed and implemented the methylation sequencing bioinformatic analysis pipeline. W.J.Z. performed the conservation analysis of methylation data between mouse and human. L.Y.C. and Y.Z. designed and implemented the methylation capture panel oligo design pipeline. L.Y.C. designed the methylation capture panel. L.L.L. produced and quality-controlled the methylation capture panel assay. Y.Z. designed and implemented the ATAC-seq and histone modification bioinformatic analysis pipelines. Y.Z. and W.J.Z. performed the single cell bioinformatic analysis. Y.Z. performed the TF footprint analysis. Y.Z., L.Y.C., J.S.L. and W.J.Z. built the statistical model for cell-free DNA methylation sequencing and performed analysis for the clinical validation test. Y.Z. and W.J wrote the paper with input from all authors.

## Acknowledgements

The authors would like to thank Dr. Yi Rao for comments on the manuscript. This study is funded by National Key R&D Program of China (2016YFC1000700, 2016YFC1000703 to A.H.Y. and 2016YFC1000400 to Y.W.), Department of Science and Technology of Guangdong Province (2016A030313787 to A.H.Y), and research fund from Euler Technology, Beijing, China.

## Conflict of interest declaration

Provisional patents were filed for the single-stranded NGS library preparation method *Tequila 7N* (WO 2020/073748) and the method for using cell-free DNA methylation pattern to predict placenta development status and pregnancy outcome (202010084924.X).

## Notes

### Summary of Updates

1. Author updated; 2. Figure 1, 2, 3, 4, 5, 6 revised; 3. Supplementary Figures revised and added; 4. Supplementary Text revised; 5. Methods updated; 6. Reference list updated

## Reference

1. Zhu, P. et al. Single-cell DNA methylome sequencing of human preimplantation embryos. Nat. Genet. 50, 12–19 (2018).

2. Xia, W. et al. Resetting histone modifications during human parental-to-zygotic transition. Science. 365, 353–360 (2019).

3. Zhou, F. et al. Reconstituting the transcriptome and DNA methylome landscapes of human implantation. Nature 572, 660–664 (2019).

4. Wu, J. et al. Chromatin analysis in human early development reveals epigenetic transition during ZGA. Nature 557, 256–260 (2018).

5. Gao, L. et al. Chromatin Accessibility Landscape in Human Early Embryos and Its Association with Evolution. Cell 173, 248-259.e15 (2018).

6. Liu, L. et al. An integrated chromatin accessibility and transcriptome landscape of human pre-implantation embryos. Nat. Commun. 10, 1–11 (2019).

7. Ziller, M. J. et al. Charting a dynamic DNA methylation landscape of the human genome. Nature 500, 477–481 (2013).

8. Guo, H. et al. The DNA methylation landscape of human early embryos. Nature 511, 606–610 (2014).

9. Lu, F. et al. Establishing chromatin regulatory landscape during mouse preimplantation development. Cell 165, 1375–1388 (2016).

10. De Iaco, A. et al. DUX-family transcription factors regulate zygotic genome activation in placental mammals. Nat. Genet. 49, 941–945 (2017).

11. Ernst, J. & Kellis, M. ChromHMM: Automating chromatin-state discovery and characterization. Nat. Methods 9, 215–216 (2012).

12. Ernst, J. & Kellis, M. Chromatin-state discovery and genome annotation with ChromHMM. Nat. Protoc. 12, 2478–2492 (2017).

13. Chen, Z., Yin, Q., Inoue, A., Zhang, C. & Zhang, Y. Allelic H3K27me3 to allelic DNA methylation switch maintains noncanonical imprinting in extraembryonic cells. Sci. Adv. 5, (2019).

14. van Heeringen, S. & Akkers, R. Principles of nucleation of H3K27 methylation during embryonic development. Genome Res. 24, 401–410 (2014).

15. Zheng, H. et al. Resetting Epigenetic Memory by Reprogramming of Histone Modifications in Mammals. Mol. Cell 63, 1066–1079 (2016).

16. Saxena, M. et al. Transcription factor-dependent ‘anti-repressive’ mammalian enhancers exclude H3K27me3 from extended genomic domains. Genes Dev. 31, 2391–2404 (2017).

17. Smith, Z. D. et al. Epigenetic restriction of extraembryonic lineages mirrors the somatic transition to cancer. Nature 549, 543–547 (2017).

18. Lyall, F., Robson, S. C. & Bulmer, J. N. Spiral artery remodeling and trophoblast invasion in preeclampsia and fetal growth restriction relationship to clinical outcome. Hypertension 62, 1046–1054 (2013).

19. Tomas, S. Z., Prusac, I. K., Roje, D. & Tadin, I. Trophoblast apoptosis in placentas from pregnancies complicated by preeclampsia. Gynecol. Obstet. Invest. 71, 250–255 (2011).

20. Maynard, S. E. et al. Excess placental soluble fms-like tyrosine kinase 1 (sFlt1) may contribute to endothelial dysfunction, hypertension, and proteinuria in preeclampsia. J. Clin. Invest. 111, 649–58 (2003).

21. Wisner, K. Gestational Hypertension and Preeclampsia. MCN Am. J. Matern. Nurs. 44, 170 (2019).

22. Vento-Tormo, R. et al. Single-cell reconstruction of the early maternal–fetal interface in humans. Nature 563, 347–353 (2018).

23. Molaro, A. et al. Sperm methylation profiles reveal features of epigenetic inheritance and evolution in primates. Cell 146, 1029–1041 (2011).

24. Gamage, T. K. J. B. et al. Human trophoblasts are primarily distinguished from somatic cells by differences in the pattern rather than the degree of global CpG methylation. Biol. Open 7, (2018).

25. Bianco-Miotto, T. et al. Recent progress towards understanding the role of DNA methylation in human placental development. Reproduction 152, R23–R30 (2016).

26. Davis, T. L. The H19 methylation imprint is erased and re-established differentially on the parental alleles during male germ cell development. Hum. Mol. Genet. 9, 2885–2894 (2000).

27. Lee, S., Ye, A. & Kim, J. DNA-binding motif of the imprinted transcription factor PEG3. PLoS One 10, 1–13 (2015).

28. Kim, J., Kollhoff, A., Bergmann, A. & Stubbs, L. Methylation-sensitive binding of transcription factor YY1 to an insulator sequence within the paternally expressed imprinted gene, Peg3. Hum. Mol. Genet. 12, 233–245 (2003).

29. Hanna, C. W. & Kelsey, G. The specification of imprints in mammals. Heredity (Edinb). 113, 176–183 (2014).

30. Jeunemaitre, X. et al. Molecular basis of human hypertension: Role of angiotensinogen. Cell 71, 169–180 (1992).

31. Zhou, A. et al. A redox switch in angiotensinogen modulates angiotensin release. Nature 468, 108–111 (2010).

32. Haase, N. et al. RNA interference therapeutics targeting angiotensinogen ameliorate preeclamptic phenotype in rodent models. J. Clin. Invest. (2020) doi: 10.1172/jci99417.

33. McGinnis, R. et al. Variants in the fetal genome near FLT1 are associated with risk of preeclampsia. Nat. Genet. 49, 1255–1260 (2017).

34. Amin-Beidokhti, M. et al. An intron variant in the FLT1 gene increases the risk of preeclampsia in Iranian women. Clin. Exp. Hypertens. 41, 697–701 (2019).

35. Low, P. & Variants, F. Protective low frequency variants for preeclampsia in the FLT1 gene in the Finnish population. Hypertension. 70, 365–371 (2018).

36. Ashar-Patel, A. et al. FLT1 and transcriptome-wide polyadenylation site (PAS) analysis in preeclampsia. Sci. Rep. 7, 1–14 (2017).

37. Thomas, C. P., Andrews, J. I. & Liu, K. Z. Intronic polyadenylation signal sequences and alternate splicing generate human soluble Fltl variants and regulate the abundance of soluble Flt1 in the placenta. FASEB J. 21, 3885–3895 (2007).

38. Fan, X. et al. Endometrial VEGF induces placental sFLT1 and leads to pregnancy complications. J. Clin. Invest. 124, 4941–4952 (2014).

39. Macintire, K. et al. PAPPA2 is increased in severe early onset pre-eclampsia and upregulated with hypoxia. Reprod. Fertil. Dev. 26, 351–357 (2014).

40. Wang, H. Y., Zhang, Z. & Yu, S. Expression of PAPPA2 in human fetomaternal interface and involvement in trophoblast invasion and migration. Genet. Mol. Res. 15, 1–16 (2016).

41. Shore, V. H. et al. Vascular endothelial growth factor, placenta growth factor and their receptors in isolated human trophoblast. Placenta 18, 657–665 (1997).

42. Cartwright, J. E. & Williams, P. J. Altered placental expression of kisspeptin and its receptor in pre-eclampsia. J. Endocrinol. 214, 79–85 (2012).

43. Zeisler, H. et al. Predictive value of the sFlt-1:PlGF ratio in women with suspected preeclampsia. N. Engl. J. Med. 374, 13–22 (2016).

44. Stewart, C. L., Kaspart, P. & Brunet, L. J. Blastocyst Implantation Depends on Maternal Expression of Leukaemia Inhibitory Factor. Nature 2664, 265–268 (1992).

45. Adelman, D. M., Gertsenstein, M., Nagy, A., Simon, M. C. & Maltepe, E. Placental cell fates are regulated in vivo by HIF-mediated hypoxia responses. Genes Dev. 14, 3191–3203 (2000).

46. Corces, M. R. et al. The chromatin accessibility landscape of primary human cancers. Science. 362, (2018).

47. Li, Z. & Schulz, M. Identification of transcription factor binding sites using Gaussian mixture models. Genome Biol. 31, 70–80 (2014).

48. Sturtevant, A. H. Studies on the bristle pattern of Drosophila. Dev. Biol. 21, 48–61 (1970).

49. Lewis, E. B. A gene complex controlling segmentation in Drosophila. Nature 276, 565–570 (1978).

50. Denell, R. E. & Frederick, R. D. Homoeosis in Drosophila: A description of the polycomb lethal syndrome. Dev. Biol. 97, 34–47 (1983).

51. Hashimoto, C., Kim, D. R., Weiss, L. A., Miller, J. W. & Morisato, D. Mutations affecting the pattern of the larval cuticle inDrosophila melanogaster_J: I. Zygotic loci on the second chromosome. Wilehm Roux Arch Dev Biol. 193, 167–282 (1984).

52. Kassis, J. A., Kennison, J. A. & Tamkun, J. W. Polycomb and Trithorax group genes in Drosophila. Genetics 206, 1699–1725 (2017).

53. Golbabapour, S. et al. Gene silencing and polycomb group proteins: An overview of their structure, mechanisms and phylogenetics. Omi. A J. Integr. Biol. 17, 283–296 (2013).

54. Abed, Jumana; Jones, R. H3K36me3 key to Polycomb-mediated gene silencing in lineage specification. Nat. Struct. Mol. Biol. 19, 1214–1215 (2012).

55. Ballaré, C. et al. Phf19 links methylated Lys36 of histone H3 to regulation of Polycomb activity. Nat. Struct. Mol. Biol. 19, 1257–1265 (2012).

56. Oguro, H. et al. Poised Lineage Specification in Multipotential Hematopoietic Stem and Progenitor Cells by the Polycomb Protein Bmi1. Cell Stem Cell 6, 279–286 (2010).

57. Ngan, C. Y. et al. Chromatin interaction analyses elucidate the roles of PRC2-bound silencers in mouse development. Nat. Genet. 52, 264–272 (2020).

58. Inoue, A., Jiang, L., Lu, F., Suzuki, T. & Zhang, Y. Maternal H3K27me3 controls DNA methylation-independent imprinting. Nature 547, 419–424 (2017).

59. Kuzmin, A. et al. The PcG gene Sfmbt2 is paternally expressed in extraembryonic tissues. Gene Expr. Patterns 8, 107–116 (2008).

60. Riising, E. M. et al. Gene silencing triggers polycomb repressive complex 2 recruitment to CpG Islands genome wide. Mol. Cell 55, 347–360 (2014).

61. Cao, Ru; Wang, Liangjun; Wang, Hengbin; Xia, Li; Erdjument-Bromage, Hediye; Tempst, Paul; Jones, Richard; Zhang, Y. Role of Histone H3 Lysine 27 Methylation in Polycomb Group Silencing. Science. 298, 1039–1043 (2002).

62. Mahadevan, S. et al. NLRP7 affects trophoblast lineage differentiation, binds to overexpressed YY1 and alters CpG methylation. Hum. Mol. Genet. 23, 706–716 (2014).

63. Inoue, K. et al. The Rodent-Specific MicroRNA Cluster within the Sfmbt2 Gene Is Imprinted and Essential for Placental Development. Cell Rep. 19, 949–956 (2017).

64. Inoue, K. et al. Loss of H3K27me3 imprinting in the Sfmbt2 miRNA cluster causes enlargement of cloned mouse placentas. Nat. Commun. 11, 1–12 (2020).

65. Turco, M. Y. et al. Trophoblast organoids as a model for maternal–fetal interactions during human placentation. Nature 564, 263–281 (2018).

66. Lyall, F., Bulmer, J. N., Kelly, H., Duffie, E. & Robson, S. C. Human trophoblast invasion and spiral artery transformation. The role of nitric oxide. Am. J. Pathol. 154, 1105–1114 (1999).

67. Qiu, X. et al. Reversed graph embedding resolves complex single-cell trajectories. Nat. Methods 14, 979–982 (2017).

68. Garrido-Gomez, T. et al. Severe pre-eclampsia is associated with alterations in cytotrophoblasts of the smooth chorion. Dev. 144, 767–777 (2017).

69. Tian, F. J. et al. The YY1/MMP2 axis promotes trophoblast invasion at the maternal-fetal interface. J. Pathol. 239, 36–47 (2016).

70. Nizyaeva, N. V. et al. Ultrastructural and Immunohistochemical Features of Telocytes in Placental Villi in Preeclampsia. Sci. Rep. 8, 1–15 (2018).

71. Lui, Y. Y. N. et al. Predominant hematopoietic origin of cell-free dna in plasma and serum after sex-mismatched bone marrow transplantation. Clin. Chem. 48, 421–427 (2002).

72. Dennis Lo, Y. M. et al. Presence of fetal DNA in maternal plasma and serum. Lancet 350, 485–487 (1997).

73. Tsui, D. W. Y., Chiu, R. W. K. & Dennis Lo, Y. M. Epigenetic approaches for the detection of fetal DNA in maternal plasma. Chimerism 1, 30–35 (2010).

74. Tong, Y. K. et al. Noninvasive prenatal detection of trisomy 21 by an epigenetic-genetic chromosome-dosage approach. Clin. Chem. 56, 90–98 (2010).

