## Supplementary Figure SF1-SF19 for "Developmentally Delayed Epigenetic Reprogramming Underlying the Pathogenesis of Preeclampsia"

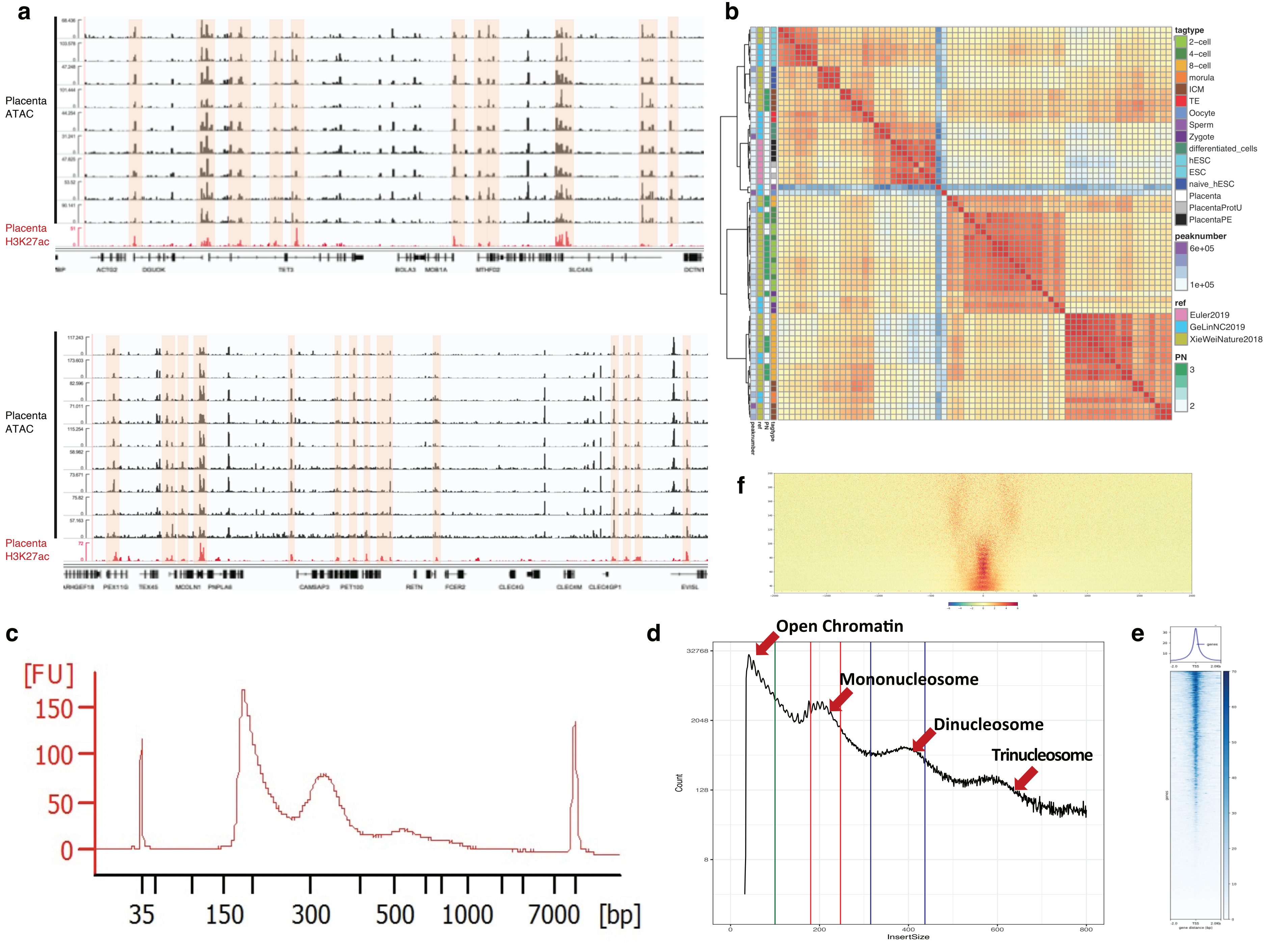


Extended Data Figure 1

#### Extended Data Figure 2

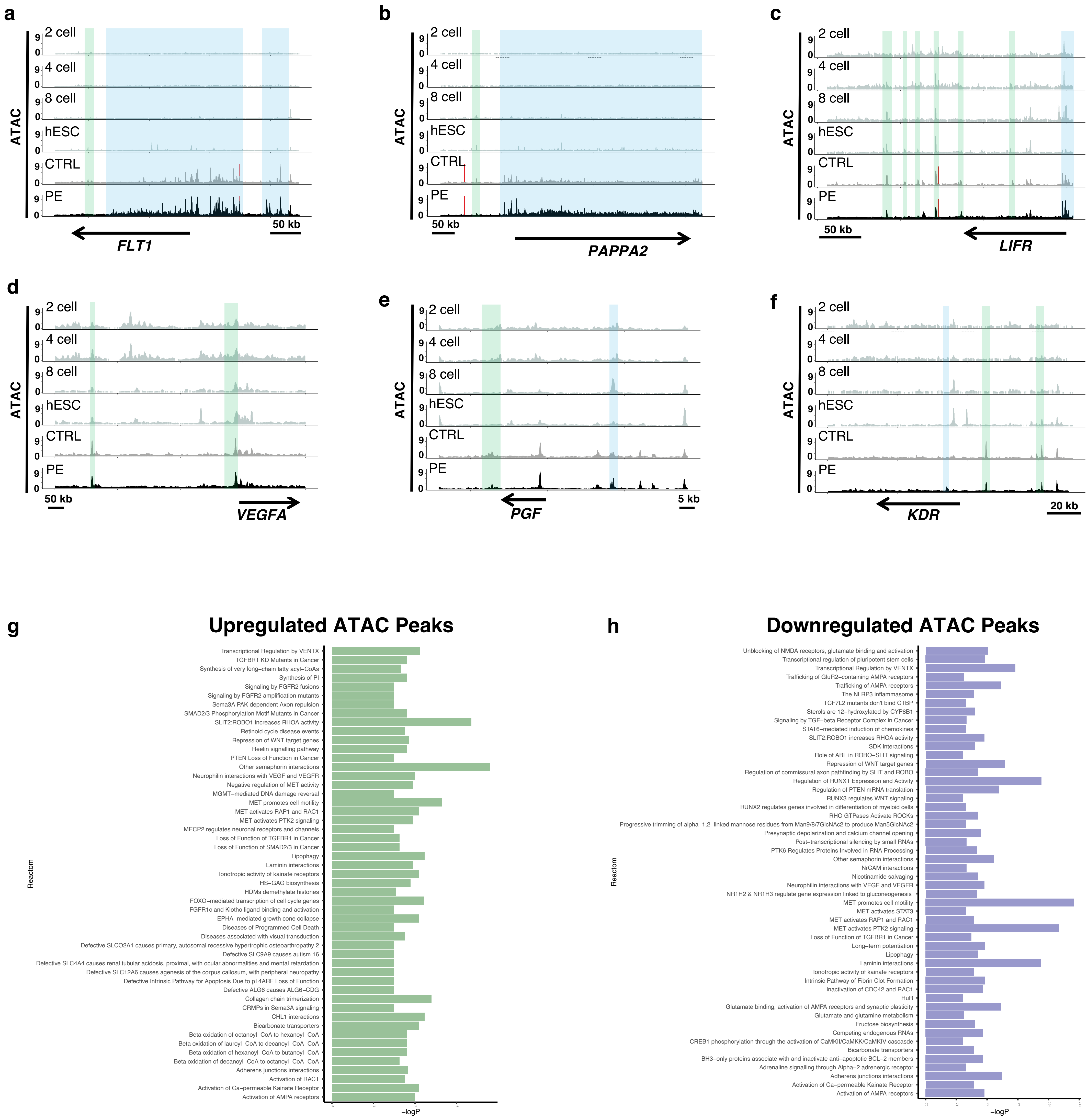

### Extended Data Figure 3

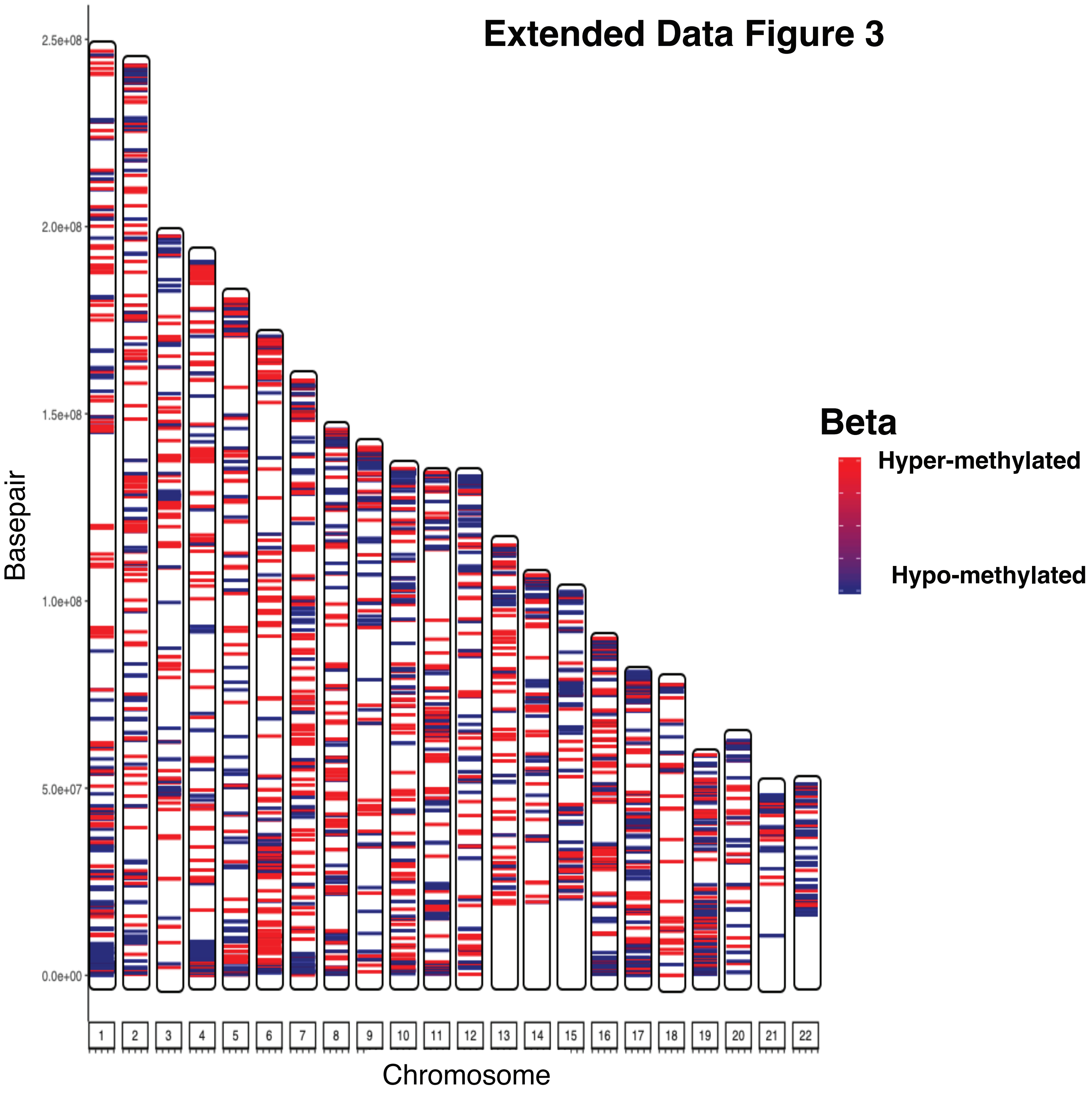

### Extended Data Figure 4

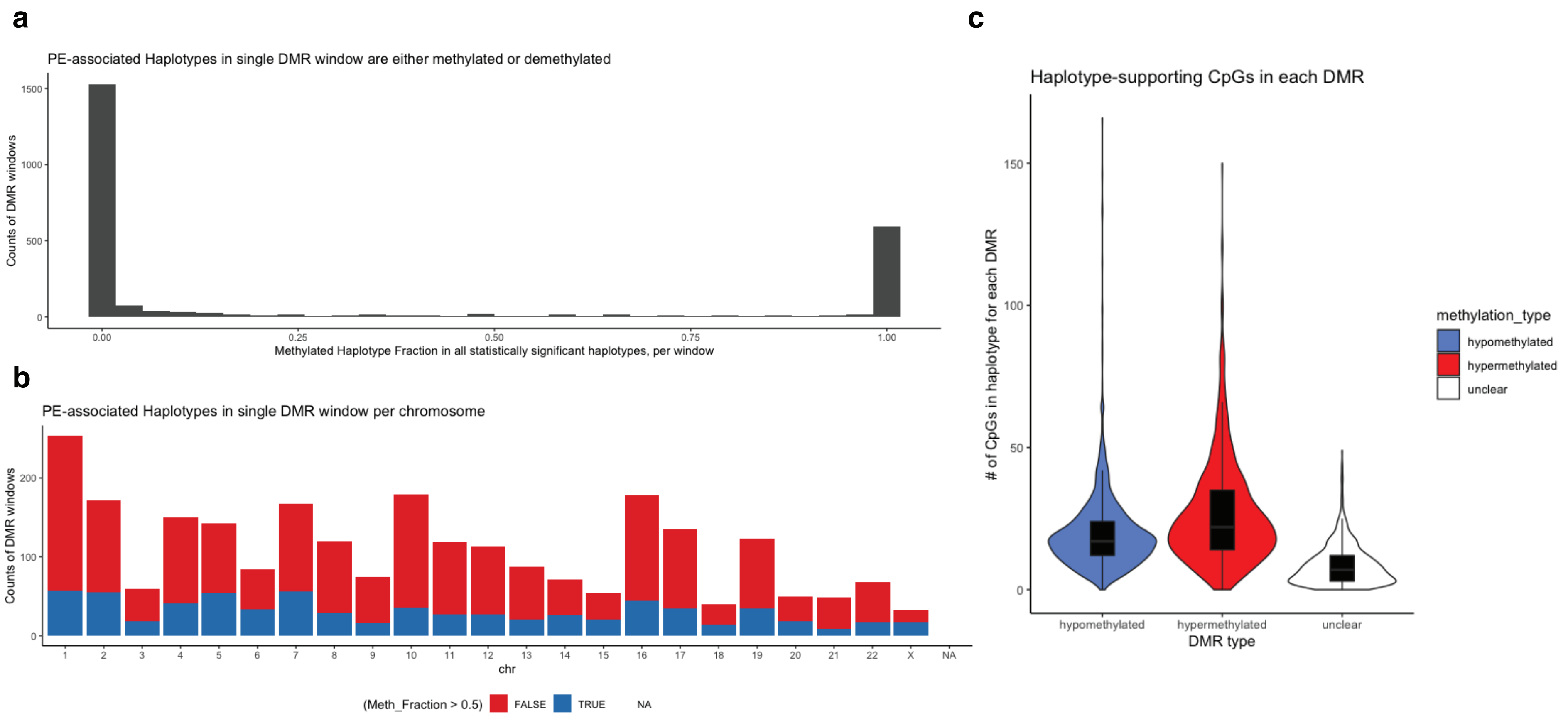

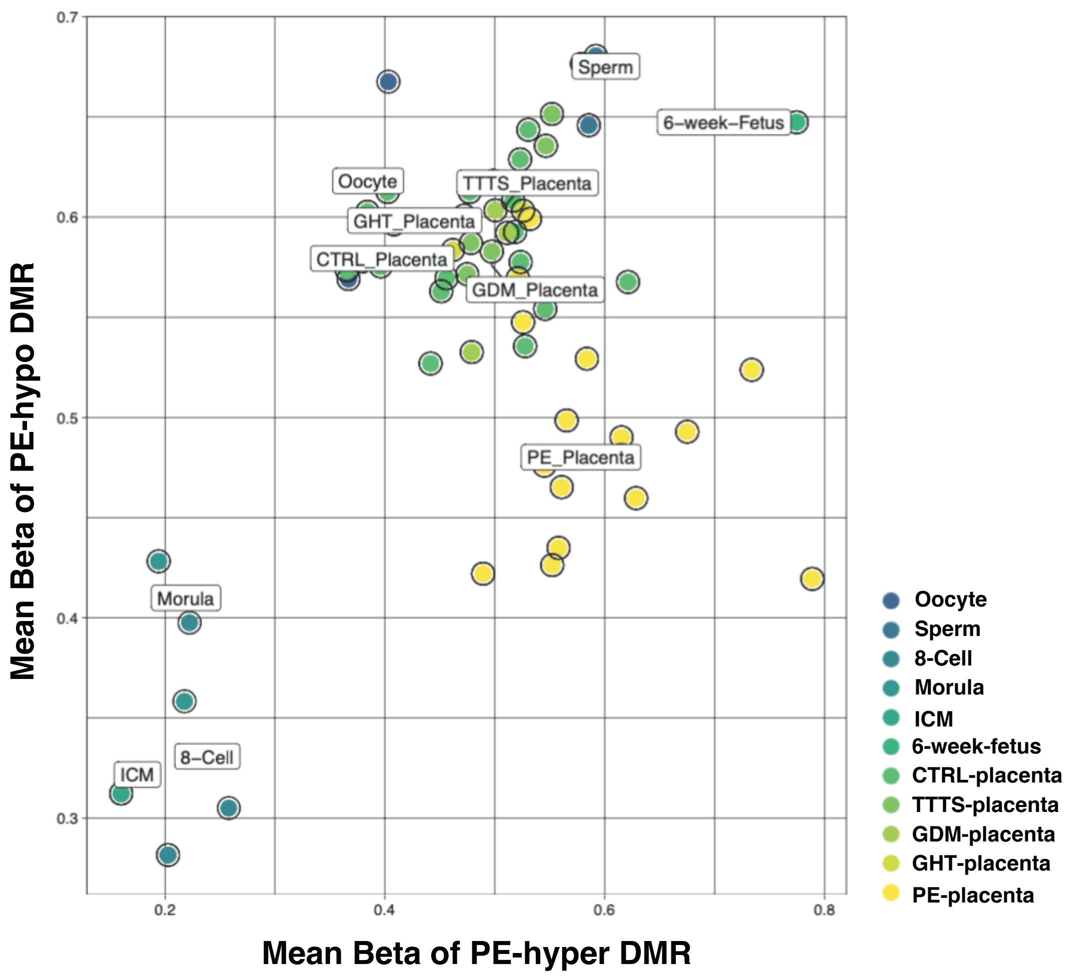


### Extended Data Figure 5

### Extended Data Figure 6

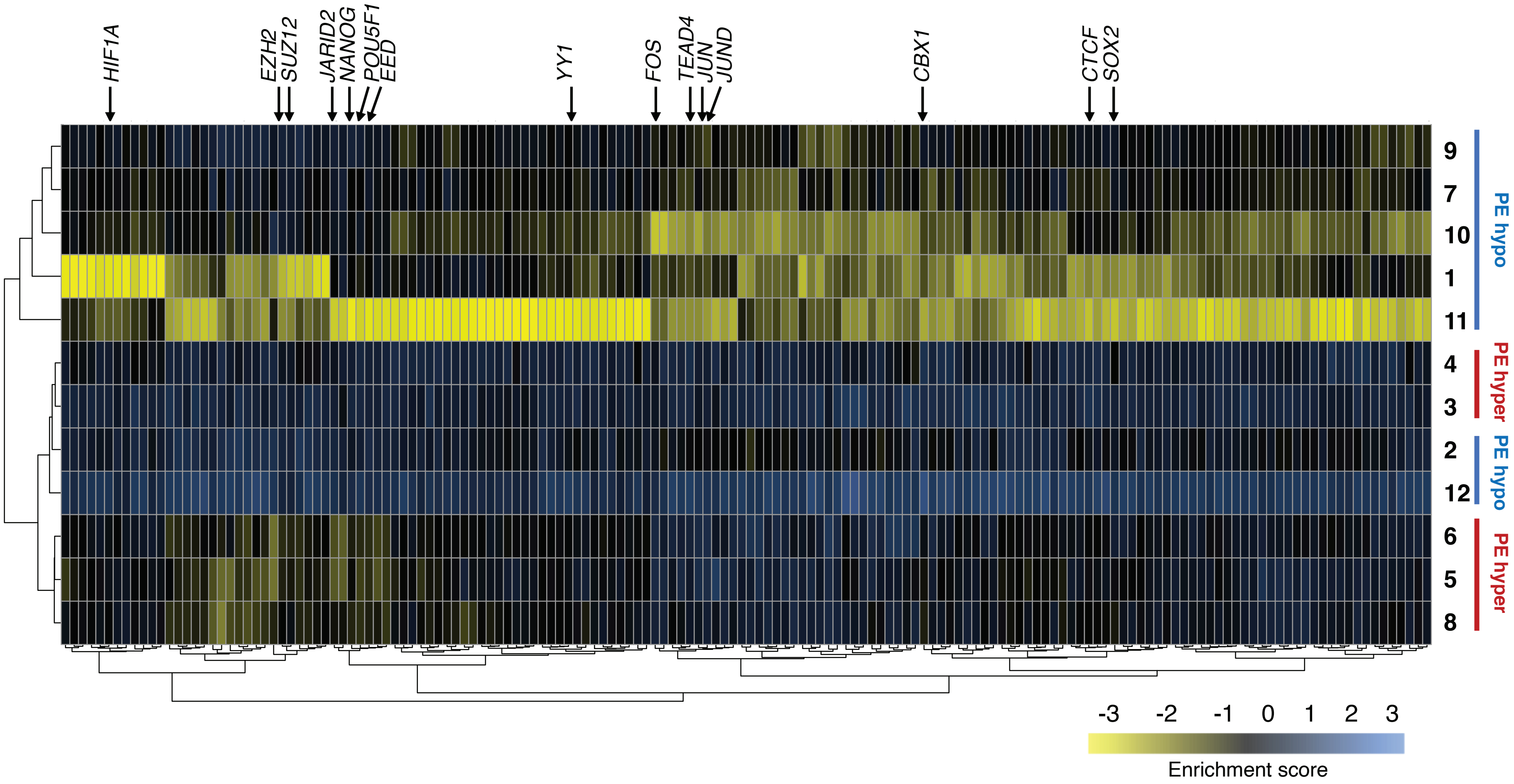

### Extended Data Figure 7

## a

##### Group6

| ID | name | Binom_Genome_Fraction | Binom_Expected | Binom_Observed_Region_Hits | Binom_Fold_Enrichment | Binom_Region_Set_Coverage | Binom_Raw_PValue | Binom_Adjp_BH | Hyper_Total_Genes | Hyper_Expected | Hyper_Observed_Gene_Hits | Hyper_Fold_Enrichment | Hyper_Gene_Set_Coverage | Hyper_Term_Gene_Coverage | Hyper_Raw_PValue | Hyper_Adjp_BH |
|---|---|---|---|---|---|---|---|---|---|---|---|---|---|---|---|---|
| GO:0030203 | glycosaminoglycan metabolic process | 0.02465235 | 6.779396 | 14 | 2.065081 | 0.05090909 | 0.00908338 | 1 | 154 | 3.229608 | 13 | 4.025256 | 0.03341902 | 0.08441558 | 2.2535E-05 | 0.02962173 |
| GO:0006022 | aminoglycan metabolic process | 0.02486353 | 6.837471 | 14 | 2.047541 | 0.05090909 | 0.00973138 | 1 | 161 | 3.376408 | 13 | 3.850245 | 0.03341902 | 0.08074534 | 3.6032E-05 | 0.03946963 |
| GO:0009952 | anterior/posterior pattern specification | 0.02773109 | 7.626051 | 14 | 1.835813 | 0.05090909 | 0.02263281 | 1 | 197 | 4.131382 | 15 | 3.630747 | 0.03856041 | 0.07614213 | 1.848E-05 | 0.02699094 |
| GO:0048568 | embryonic organ development | 0.0656595 | 18.05636 | 26 | 1.439936 | 0.09454545 | 0.04039771 | 1 | 419 | 8.787051 | 24 | 2.731292 | 0.06169666 | 0.05727924 | 9.4491E-06 | 0.01774413 |
| GO:0009790 | embryo development | 0.1307873 | 35.96651 | 46 | 1.278967 | 0.1672727 | 0.04760969 | 1 | 906 | 19.00016 | 44 | 2.31577 | 0.1131105 | 0.04856512 | 1.9634E-07 | 0.00258083 |
| GO:0001501 | skeletal system development | 0.06999713 | 19.24921 | 27 | 1.402655 | 0.09818182 | 0.04852265 | 1 | 465 | 9.751739 | 27 | 2.768737 | 0.06940874 | 0.05806452 | 2.0176E-06 | 0.00552903 |

## b

##### Group11

| ID | name | Binom_Genome_Fraction | Binom_Expected | Binom_Observed_Region_Hits | Binom_Fold_Enrichment | Binom_Region_Set_Coverage | Binom_Raw_PValue | Binom_Adjp_BH | Hyper_Total_Genes | Hyper_Expected | Hyper_Observed_Gene_Hits | Hyper_Fold_Enrichment | Hyper_Gene_Set_Coverage | Hyper_Term_Gene_Coverage | Hyper_Raw_PValue | Hyper_Adjp_BH |
|---|---|---|---|---|---|---|---|---|---|---|---|---|---|---|---|---|
| GO:0007389 | pattern specification process | 0.06274844 | 14.80863 | 31 | 2.093374 | 0.1313559 | 8.7499E-05 | 0.04521248 | 406 | 8.054774 | 26 | 3.227899 | 0.07065217 | 0.06403941 | 1.7311E-07 | 0.00013814 |
| GO:0003002 | regionalization | 0.04922166 | 11.61631 | 30 | 2.582575 | 0.1271186 | 2.2687E-06 | 0.00928799 | 313 | 6.209715 | 25 | 4.02595 | 0.06793478 | 0.0798722 | 3.9776E-09 | 2.6143E-05 |
| GO:0045165 | cell fate commitment | 0.04712743 | 11.12207 | 29 | 2.607427 | 0.1228814 | 2.8263E-06 | 0.00928799 | 226 | 4.483692 | 23 | 5.129701 | 0.0625 | 0.1017699 | 1.5231E-10 | 2.0022E-06 |
| GO:0061448 | connective tissue development | 0.0328432 | 7.750995 | 21 | 2.70933 | 0.08898305 | 4.1913E-05 | 0.02899694 | 198 | 3.92819 | 16 | 4.073123 | 0.04347826 | 0.08080808 | 2.2504E-06 | 0.00123257 |
| GO:0009952 | anterior/posterior pattern specification | 0.02773109 | 6.544538 | 20 | 3.055983 | 0.08474576 | 1.2051E-05 | 0.01416103 | 197 | 3.908351 | 18 | 4.605523 | 0.04891304 | 0.09137056 | 8.2983E-08 | 0.00010515 |
| GO:0035270 | endocrine system development | 0.0226831 | 5.353212 | 18 | 3.362468 | 0.07627119 | 9.5338E-06 | 0.01392467 | 121 | 2.400561 | 12 | 4.998832 | 0.0326087 | 0.09917355 | 5.2225E-06 | 0.0022145 |
| GO:0051147 | regulation of muscle cell differentiation | 0.02364063 | 5.579189 | 17 | 3.047038 | 0.0720339 | 5.7082E-05 | 0.03573049 | 169 | 3.352849 | 14 | 4.175553 | 0.03804348 | 0.08284024 | 7.2695E-06 | 0.0028105 |
| GO:0051216 | cartilage development | 0.02517865 | 5.94216 | 17 | 2.860912 | 0.0720339 | 0.00012118 | 0.04977839 | 152 | 3.01558 | 13 | 4.310945 | 0.03532609 | 0.08552632 | 1.091E-05 | 0.0034979 |
| GO:0048663 | neuron fate commitment | 0.0158371 | 3.737555 | 14 | 3.745764 | 0.05932203 | 3.0302E-05 | 0.02212891 | 68 | 1.349075 | 11 | 8.153732 | 0.0298913 | 0.1617647 | 8.9999E-08 | 0.00010515 |
| GO:0021515 | cell differentiation in spinal cord | 0.00932891 | 2.201622 | 11 | 4.996317 | 0.04661017 | 1.711E-05 | 0.01711625 | 53 | 1.051485 | 8 | 7.608285 | 0.02173913 | 0.1509434 | 9.0431E-06 | 0.00329127 |
| GO:0010463 | mesenchymal cell proliferation | 0.00570746 | 1.346961 | 8 | 5.939297 | 0.03389831 | 7.5421E-05 | 0.04310482 | 17 | 0.3372689 | 5 | 14.82497 | 0.01358696 | 0.2941176 | 1.5208E-05 | 0.0039981 |
| GO:0021871 | forebrain regionalization | 0.00434386 | 1.025151 | 8 | 7.803726 | 0.03389831 | 1.1165E-05 | 0.01416103 | 24 | 0.4761443 | 7 | 14.70143 | 0.01902174 | 0.2916667 | 2.9569E-07 | 0.00019434 |
| GO:0048333 | mesodermal cell differentiation | 0.00483111 | 1.140143 | 8 | 7.016665 | 0.03389831 | 2.3697E-05 | 0.02002209 | 26 | 0.515823 | 6 | 11.6319 | 0.01630435 | 0.2307692 | 9.6328E-06 | 0.0033322 |
| GO:0021978 | telencephalon regionalization | 0.00302486 | 0.7138669 | 6 | 8.404928 | 0.02542373 | 9.5258E-05 | 0.04637661 | 13 | 0.2579115 | 5 | 19.3865 | 0.01358696 | 0.3846154 | 3.3771E-06 | 0.00164415 |
| GO:0021798 | forebrain dorsal/ventral pattern formation | 0.0010305 | 0.2431982 | 5 | 20.55936 | 0.02118644 | 5.5739E-06 | 0.01221146 | 7 | 0.1388754 | 4 | 28.8028 | 0.01086957 | 0.5714286 | 5.0887E-06 | 0.0022145 |

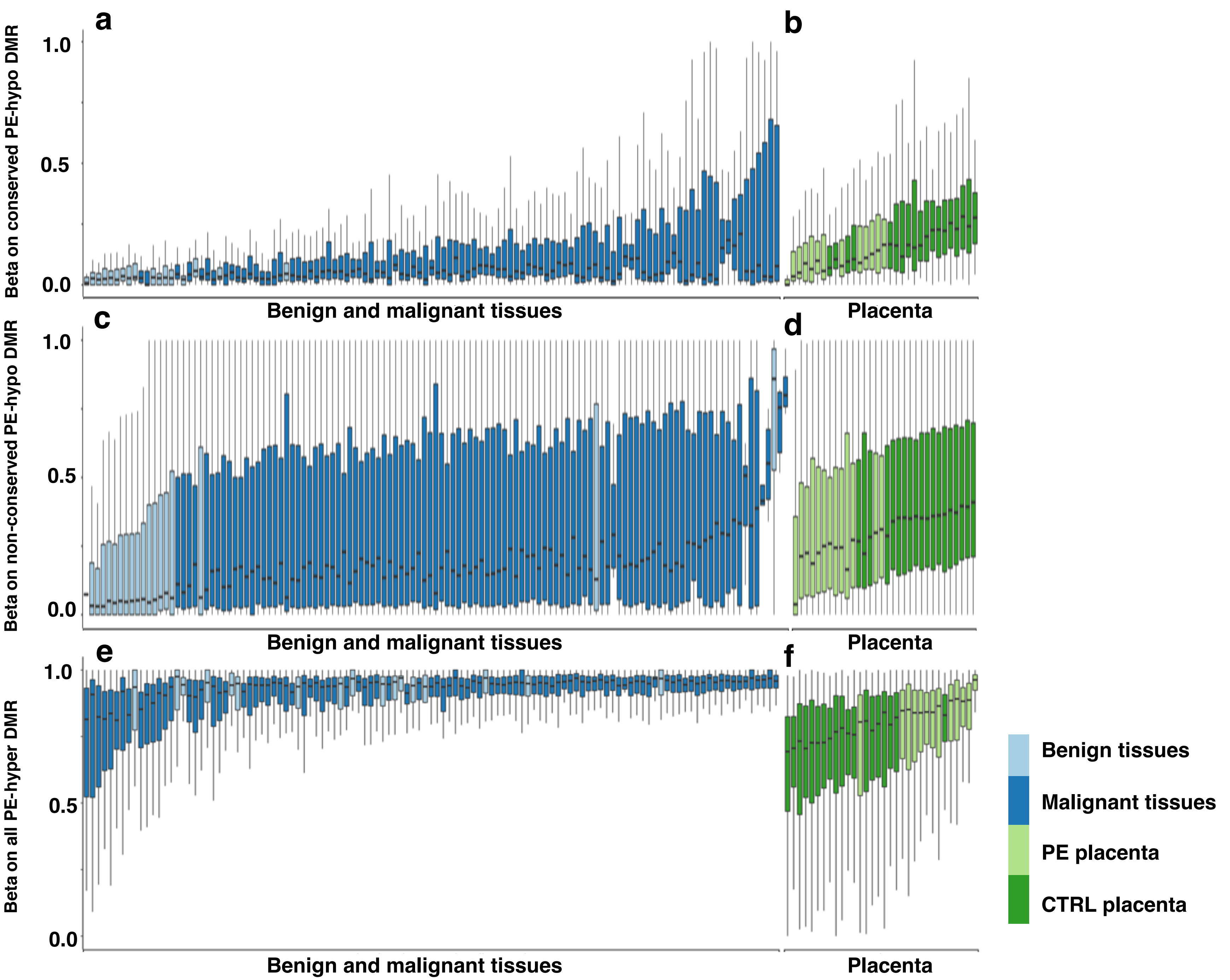


**Extended Data Figure 8**

#### Extended Data Figure 9

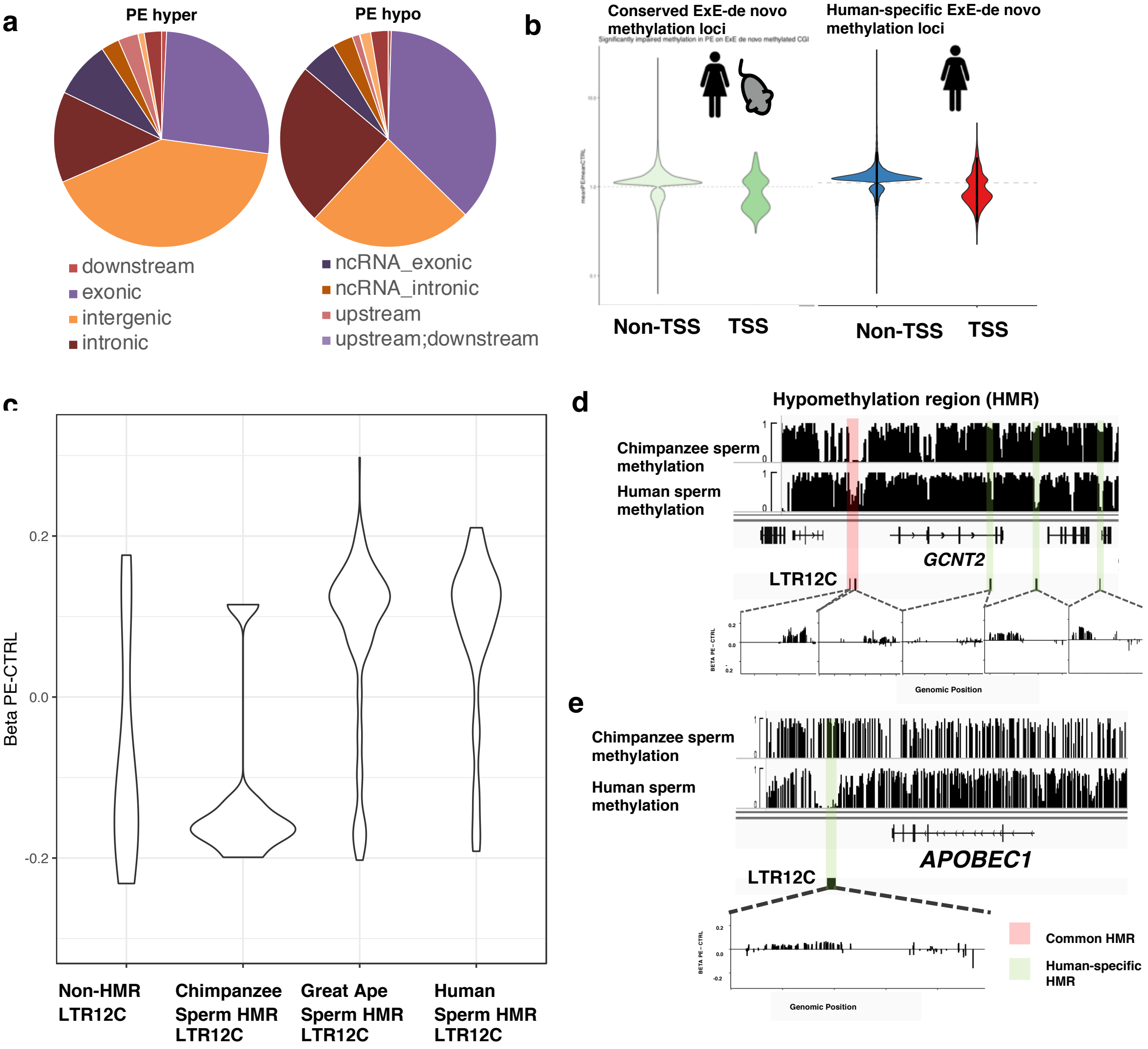

### Extended Data Figure 10

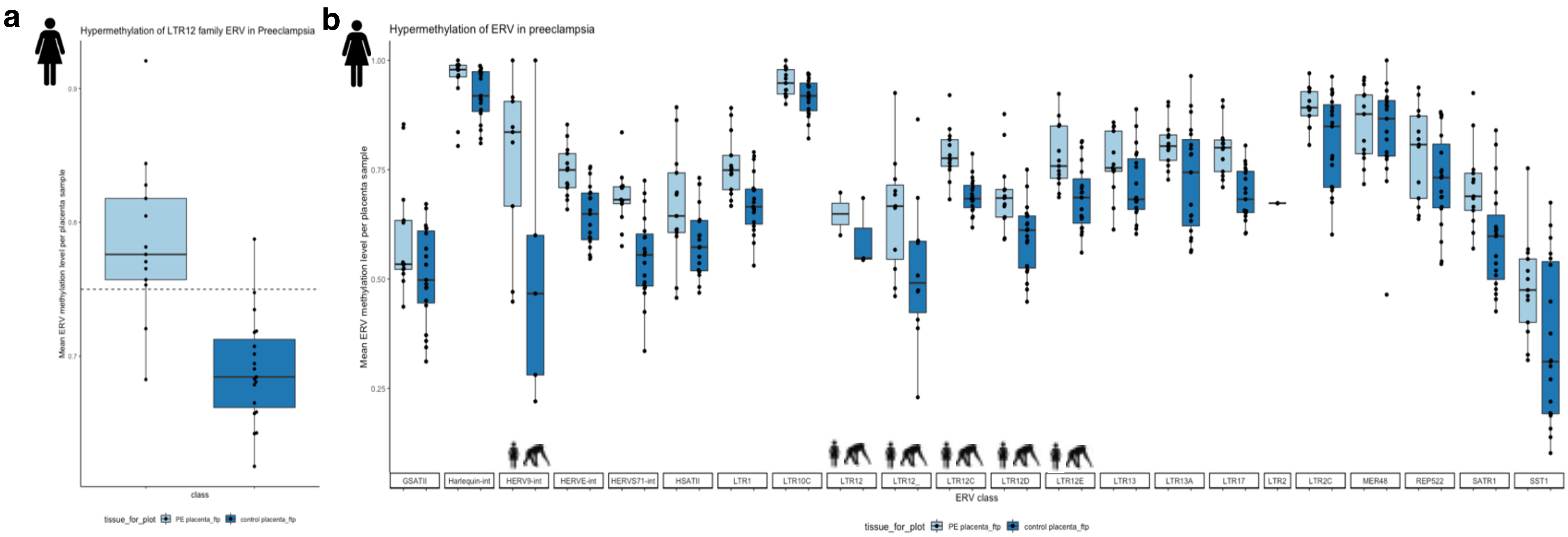

### Extended Data Figure 11

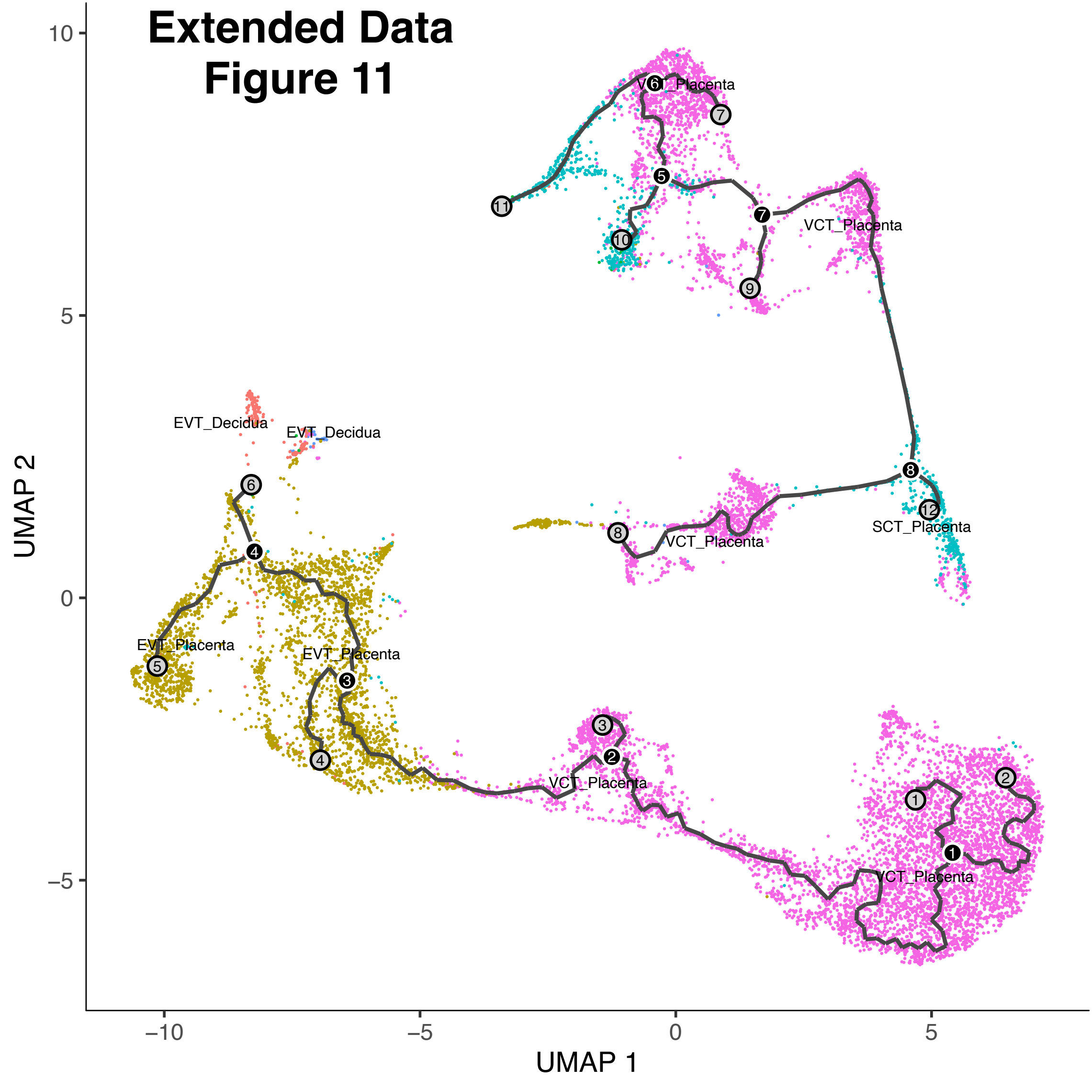

### Extended Data Figure 12

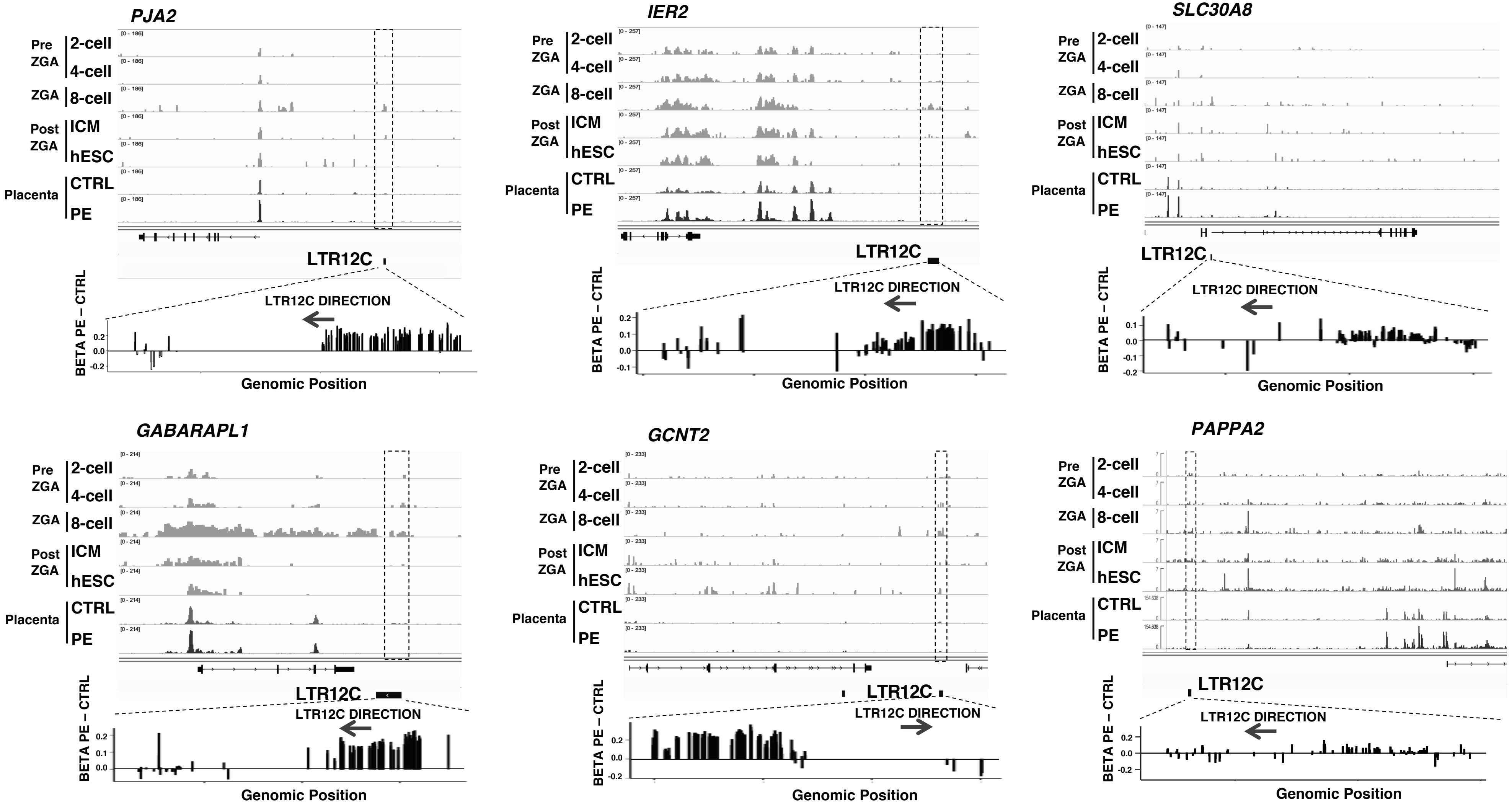

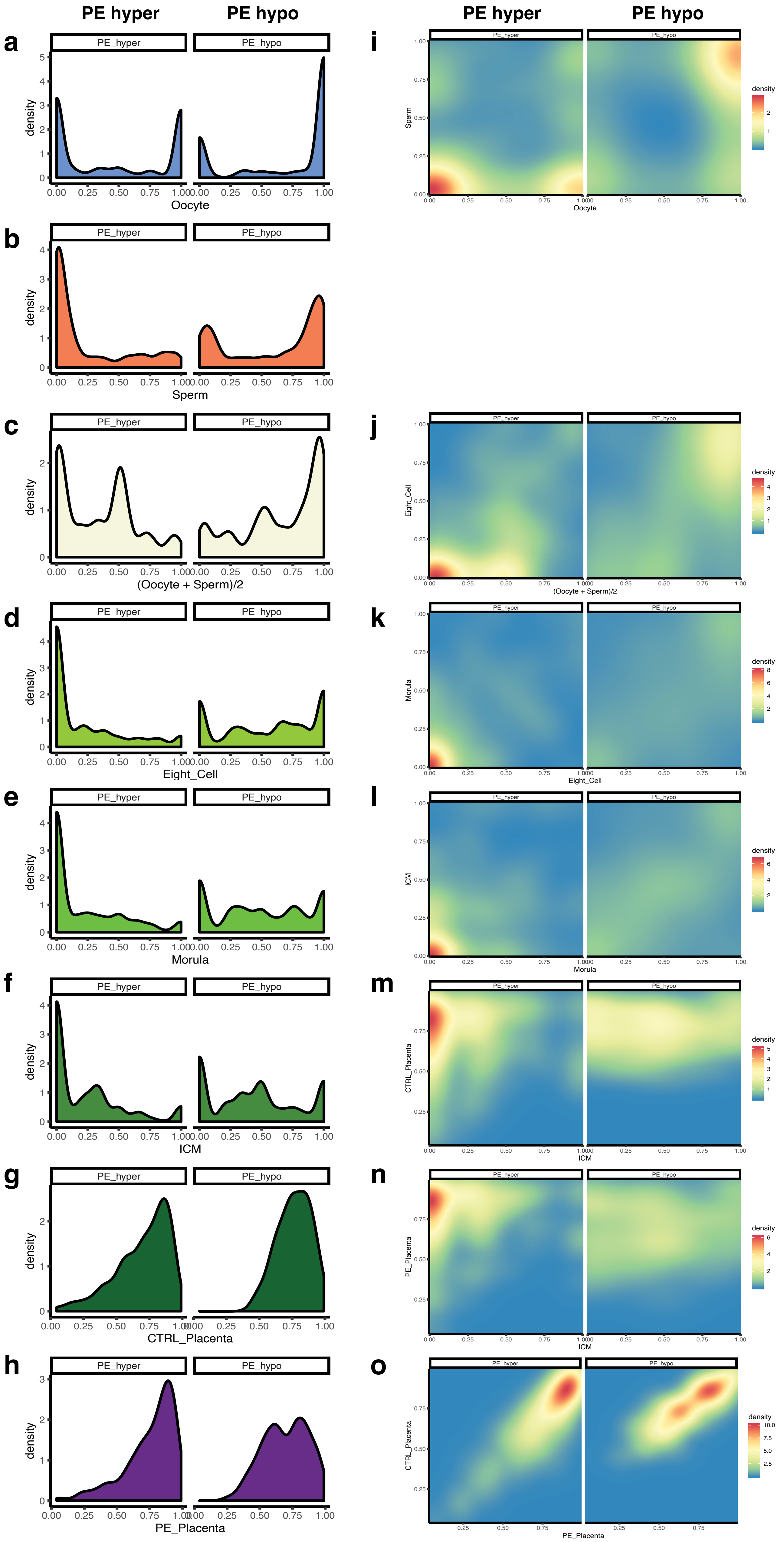


**Extended Data Figure 13**

### Extended Data Figure 14

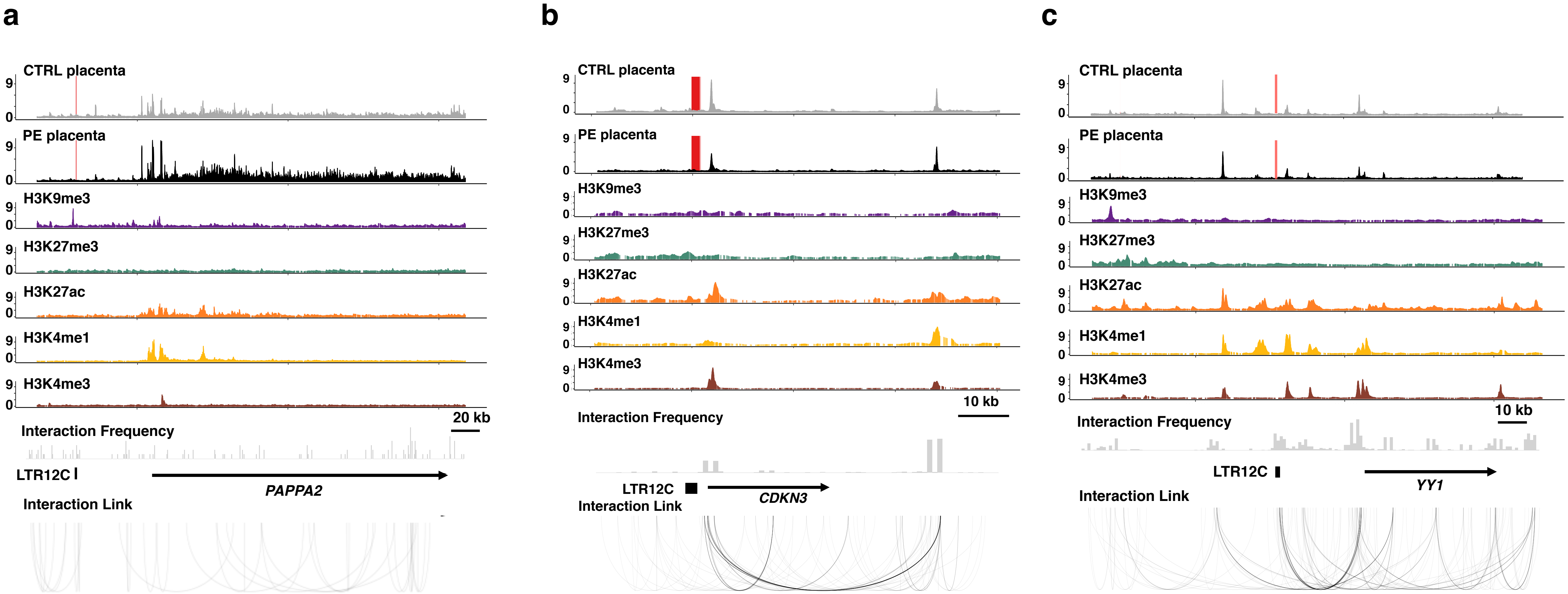

### Extended Data Figure 15

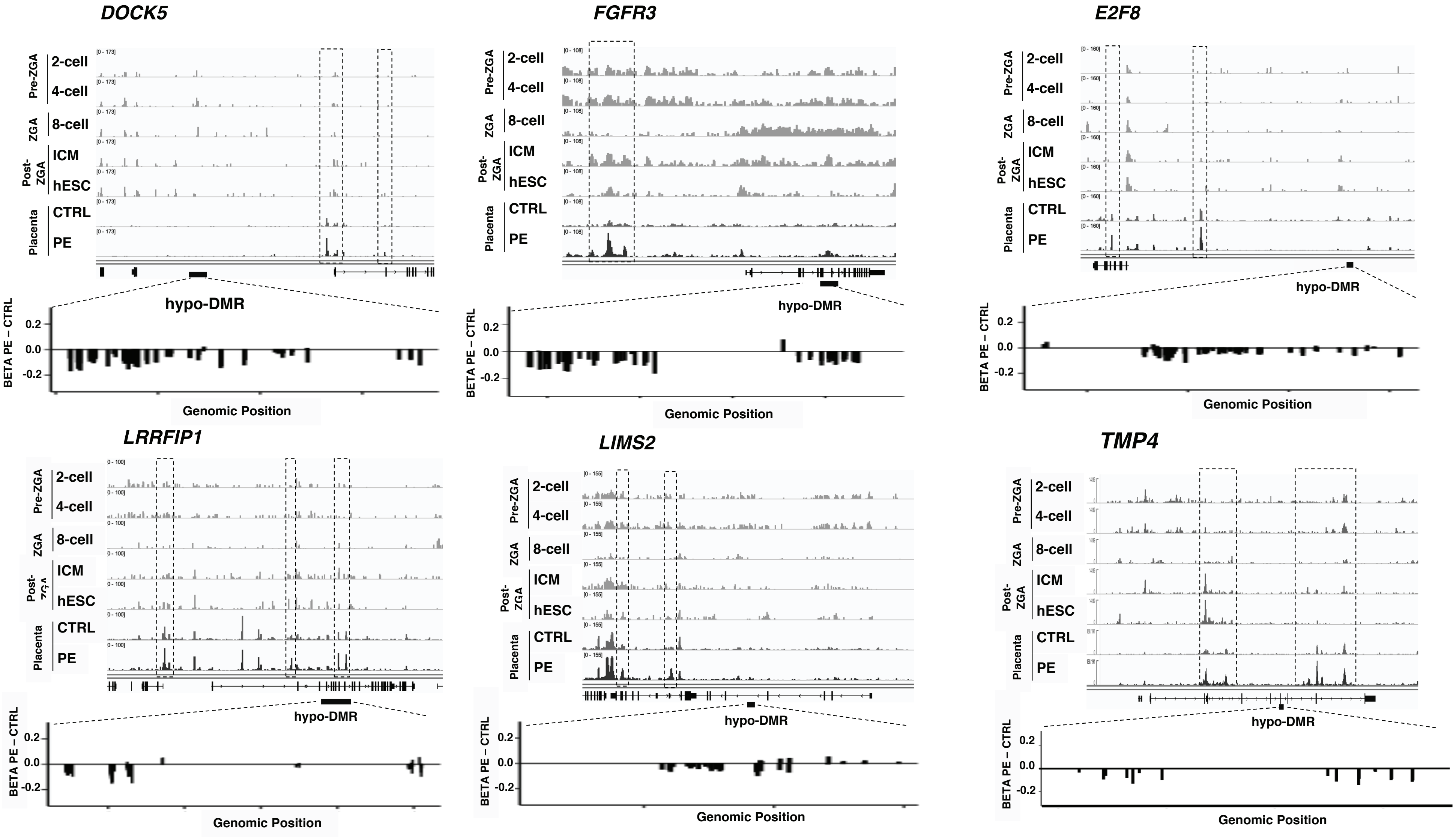

### Extended Data Figure 16

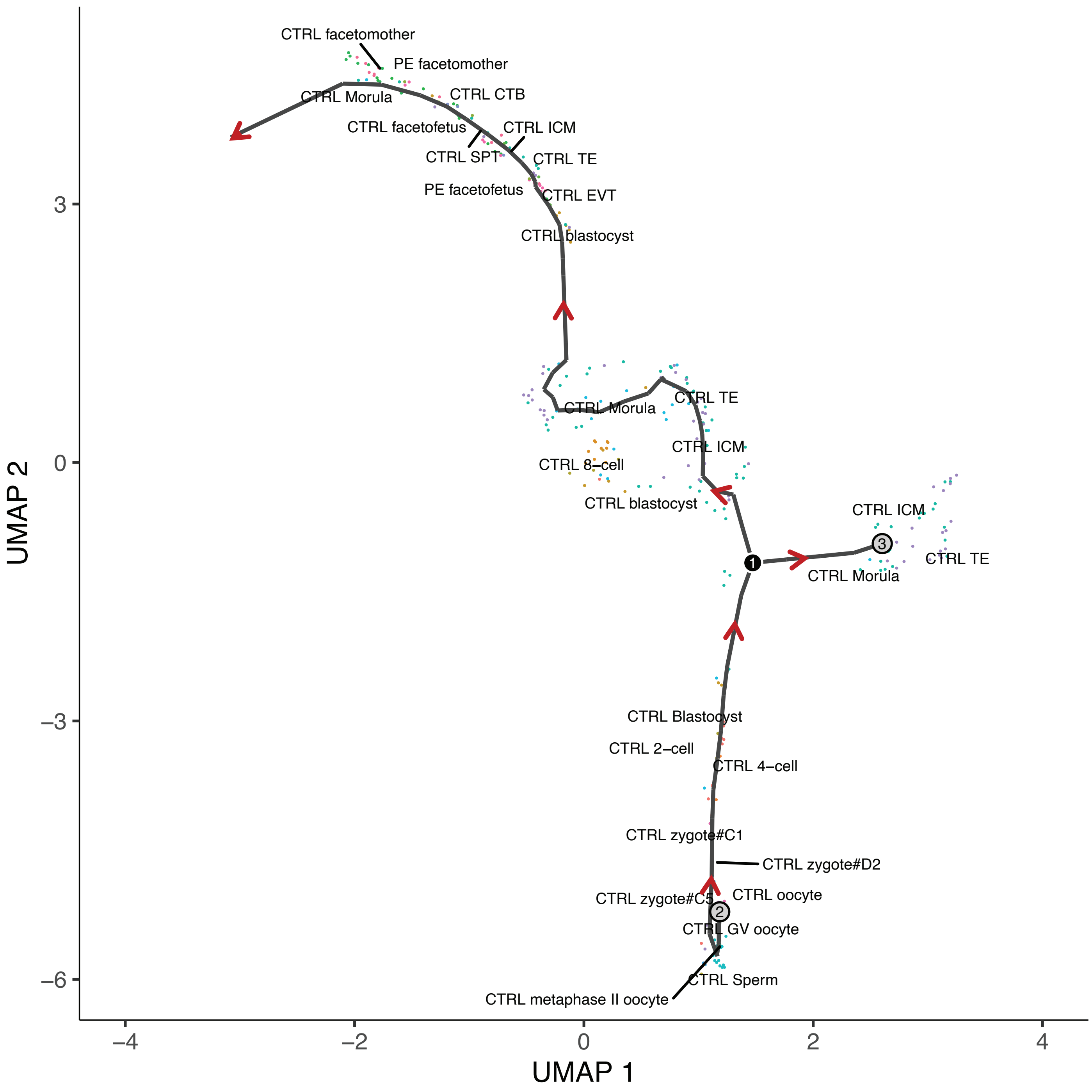

### Extended Data Figure 17

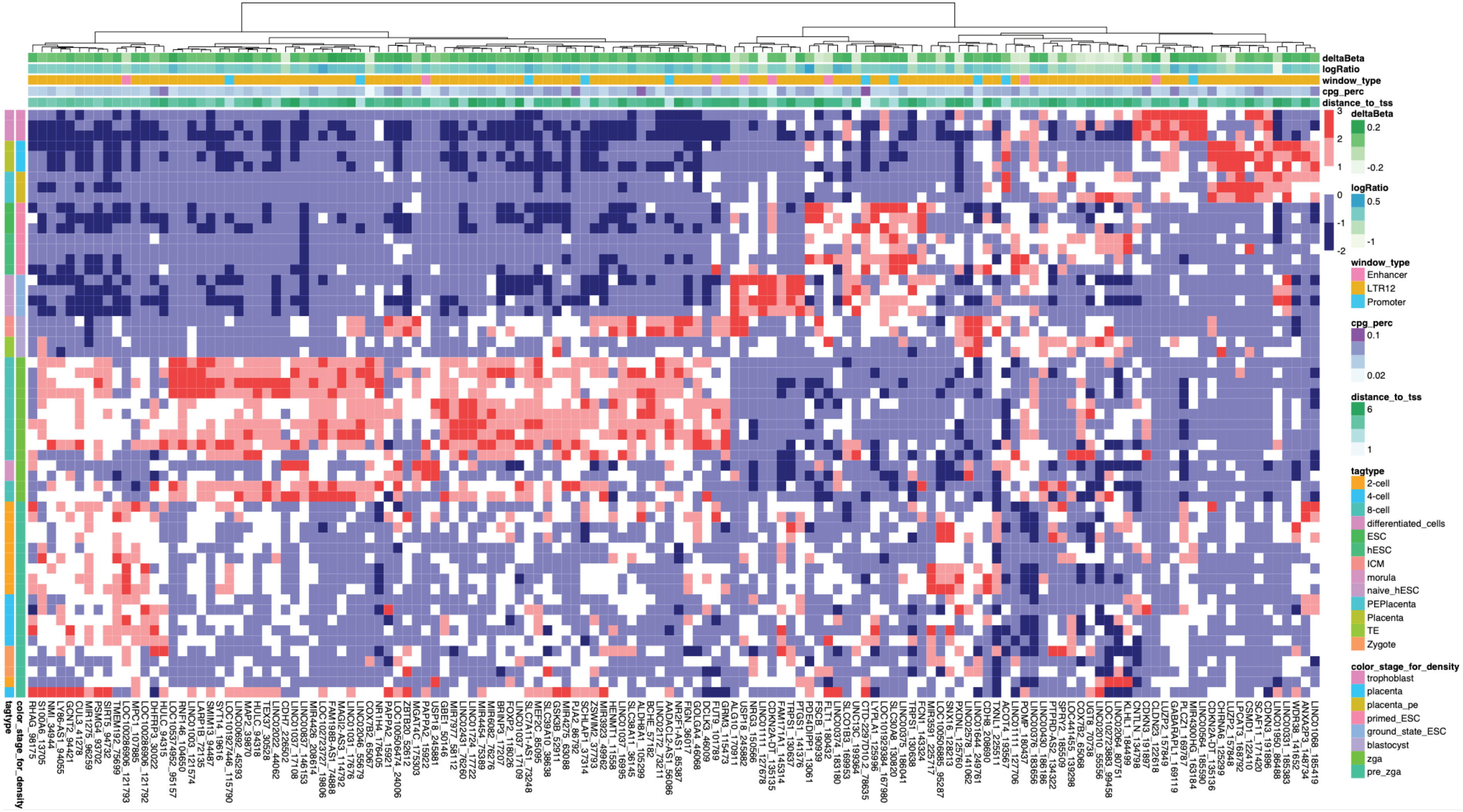

### Extended Data Figure 18

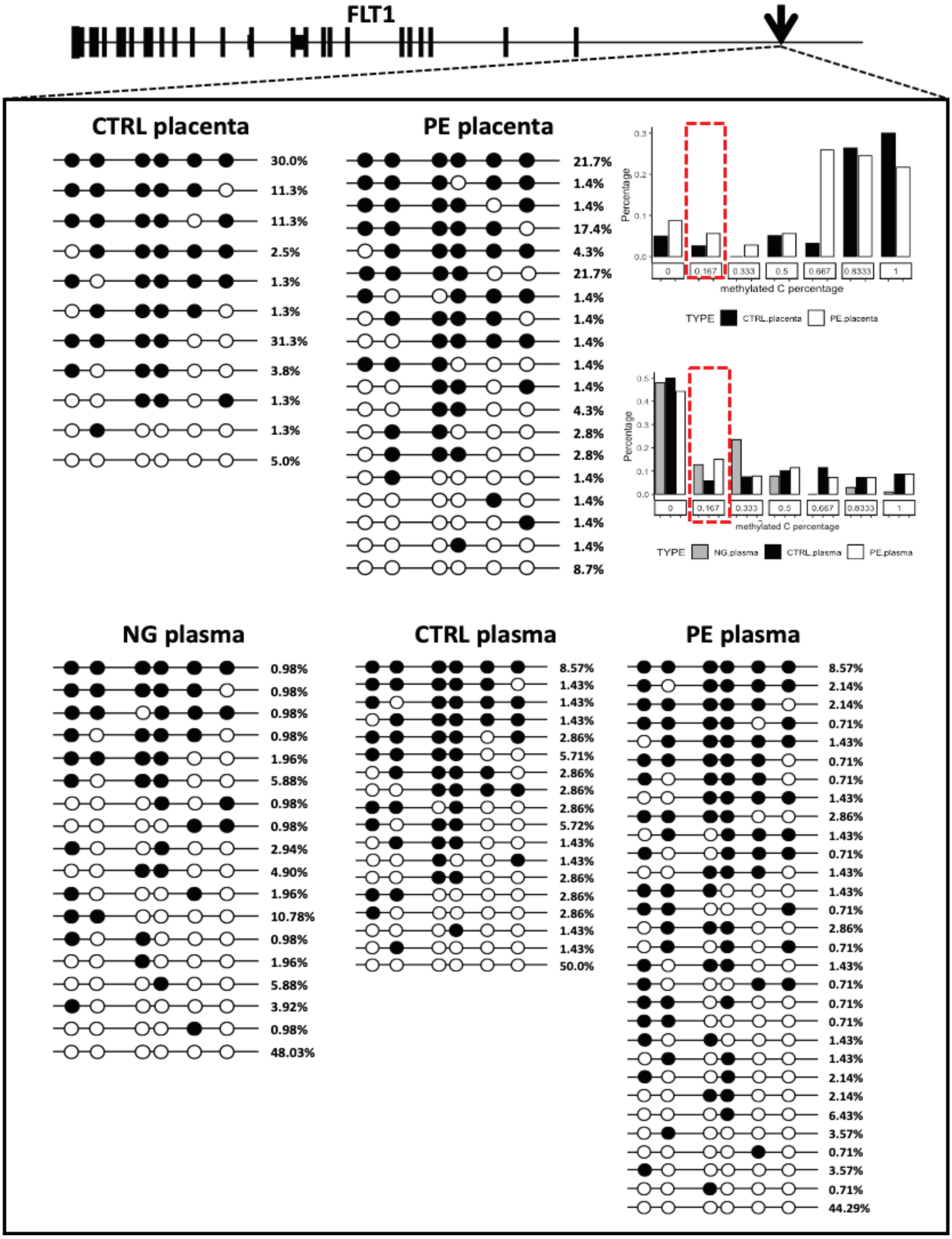

### Extended Data Figure 19

**a**

| Element Type | All | Overall ATAC Difference | Gain in Preeclampsia | Loss in Preeclampsia | Percent Gain in Preeclampsia | Percent Loss in Preeclampsia |
|---|---|---|---|---|---|---|
| Active TSS | 18422 | 192 | 108 | 84 | 0.005862556 | 0.004559765 |
| CpG island | 28691 | 170 | 91 | 79 | 0.003171726 | 0.002753477 |
| Weak Repressed PolyComb | 36018 | 189 | 113 | 76 | 0.00313732 | 0.002110056 |
| Weak transcription | 110506 | 735 | 340 | 395 | 0.003076756 | 0.003574467 |
| Transcripts at gene 5 and 3 | 679 | 5 | 2 | 3 | 0.002945508 | 0.004418262 |
| Bivalent Enhancer | 10425 | 39 | 22 | 17 | 0.002110312 | 0.001630695 |
| Flanking Bivalent TSS Enhancer | 7118 | 24 | 15 | 9 | 0.002107334 | 0.0012644 |
| Enhancers | 119230 | 477 | 248 | 229 | 0.002080013 | 0.001920658 |
| Strong transcription | 33323 | 145 | 65 | 80 | 0.001950605 | 0.002400744 |
| Flanking Active TSS | 17065 | 69 | 32 | 37 | 0.001875183 | 0.00216818 |
| Genic enhancers | 8162 | 32 | 14 | 18 | 0.001715266 | 0.002205342 |
| Bivalent Poised TSS | 3614 | 11 | 6 | 5 | 0.00166021 | 0.001383509 |
| Repressed PolyComb | 15998 | 32 | 18 | 14 | 0.001125141 | 0.000875109 |
| ZNF genes and repeats | 14193 | 19 | 14 | 5 | 0.000986402 | 0.000352286 |
| Heterochromatin | 42046 | 70 | 39 | 31 | 0.000927556 | 0.000737288 |

**b**

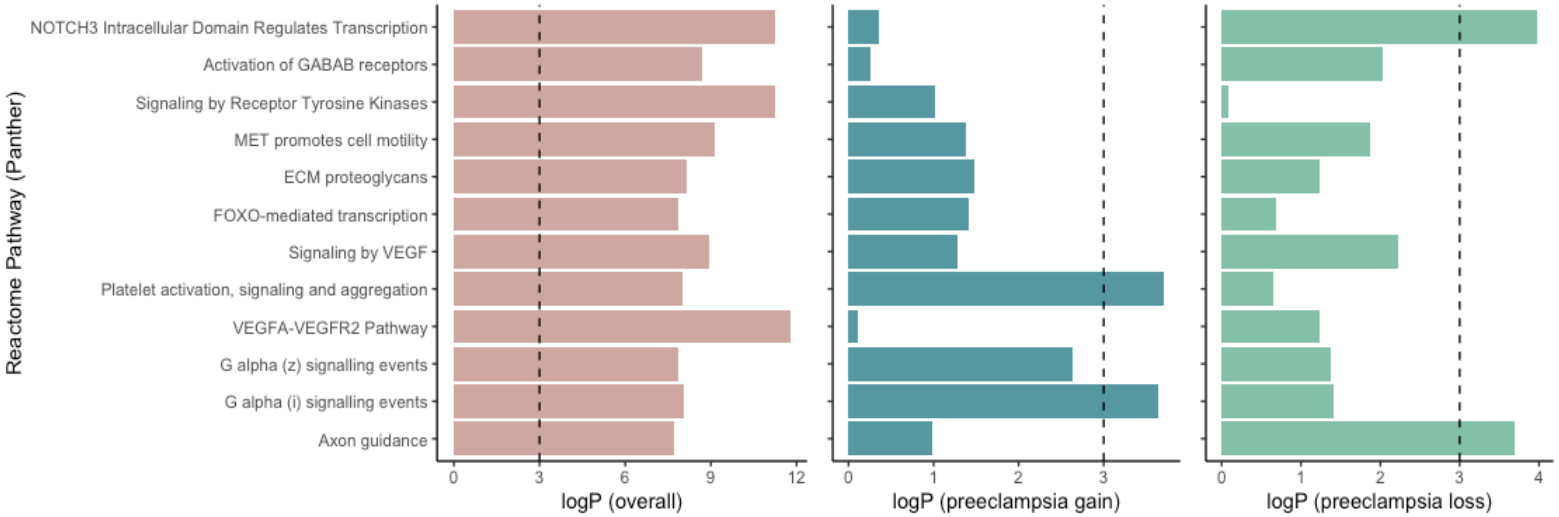
