## Supplementary Text for "Developmentally Delayed Epigenetic Reprogramming Underlying the Pathogenesis of Preeclampsia"

This file includes:

Materials and Methods
Reference List for Supplementary Materials
Legends for Extended Data Figure 1-19
Legends for Extended Data Table S1-S6

### Materials and Methods

#### Human biospecimens

This study was conducted in accordance with the measures of the People's Republic of China on the administration of Human Assisted Reproductive Technology, the ethical principles of the Human Assisted Reproductive Technology, the Helsinki declaration, and the internal ethic protocols of Guangdong Woman and Children's Hospital, Peking University Third Hospital, Zhongnan Hospital of Wuhan University, and Shanghai Changzheng Hospital.

The study protocol was approved by the Institutional Review Board (IRB) of Guangdong Woman and Children's Hospital (YiLun[201701044]), IRB of Peking University Third Hospital (IRB0001052-16009), IRB of Changzheng Hospital, Shanghai, China (2017SL016), and the Ethics Committee at Zhongnan Hospital of Wuhan University (approval number: 2015029). The human sample preservation by Department of Biological Repositories, Zhongnan Hospital of Wuhan University (official member of the International Society for Biological and Environmental Repositories-International Repository Locator, https://irlocator.isber.org/details/60) was approved by the Ethics Committee (approval number: 2017038) and China Human Genetic Resources Management Office, Ministry of Science and Technology of The People's Republic of China (approval number: 20171793).

For placenta ATAC-seq experiments, placenta samples were collected from Peking University Third Hospital. For placenta methylation capture sequencing experiments, placenta samples were collected from Guangdong Woman and Children's Hospital and Peking University Third Hospital. For biological validation tests on tumor and normal samples, FFPE tumor or normal tissue slides were collected from Zhongnan Hospital of Wuhan University, Wuhan, China and Changzheng Hospital, Shanghai, China.

To validate the predictive power of cell-free DNA methylation pattern on pregnancy outcomes (preeclampsia v.s. non-preeclampsia), we retrospectively collected archived serum samples left from non-invasive prenatal testing (NIPT) clinical tests in Guangdong Woman and Childrens' Hospital from 2015-2017. Together, we collected a cohort of 159 preeclampsia- and non-preeclampsia pregnant females, whose clinical characteristics were denoted in Extended Data Table S6. The clinical characteristics were blinded for molecular biologists and bioinformatician by the clinician. To validate the predictive power in a prospective setting, two (2) volunteers at gestational age 12-13 with a preeclampsia medical history for prospective pregnancy outcome prediction were recruited in Guangdong Woman and Childrens' Hospital at 2019. Written consent was obtained from all volunteers.

Clinical assessment of human (pregnant female) phenotype was done according to ACOG guideline of hypertension in pregnancy[1]. Routine laboratory tests and pathology assessments were done according to the relevant Chinese clinical protocols.

Human serum was collected with MiniMax cell-free DNA tube (Apostle Inc. CA) or Cell-free DNA BCT (Streck, Inc. NE) (Beijing) or BD EDTA tube (Guangdong) according to the manufacturer's protocol and stored at room temperature (cell-free DNA tube) or 4 degree Celsius (EDTA tube) for no longer than 8 hours before serum separation. Serum was separated with two consecutive low-speed centrifugation (1500x rpm) on a tabletop centrifuge. Placenta was collected within 1hr of labor and transferred to laboratory in high glucose, 10% FBS supplemented DMEM. Placenta sample was resected in PBS from the maternal or fetal surface of placenta, 2cm*1cm (diameter*height) in size and within 4-5cm from the root of umbilical cord. For ATAC-seq, villous was manually dissected from the maternal surface placenta with a pair of fine forceps for further processing. For methylation sequencing, placenta samples were flash frozen in liquid nitrogen and stored at -80 degree Celsius.

### Public data

The study used a series of human subject data from NCBI SRA sequencing read archive (https://www.ncbi.nlm.nih.gov/sra) and the Chinese Genome Sequencing Archive (https://bigd.big.ac.cn/) including human embryo ATAC-seq [2,3], human embryo methylation sequencing[4], human embryo Cut-And-Run histone modification sequencing[5], human placenta single cell RNAseq[6], human trophoblast RRBS[7], human and chimpanzee sperm methylation sequencing[8]. Human embryo DnaseI sequencing result[9] was initially used for comparison but not used in further analysis because of a failure of batch-effect removal. Placenta and BMP4-primed trophoblast H3K27ac, H3K4me3, H3K4me1, H3K9me3 and H3K27me3 ChIP-seq data were collected from Roadmap project website (http://www.roadmapepigenomics.org/data). The Encode transcription factor binding sites, ChromHMM tracks, CpG islands, repeat regions data, and lift-over chains were collected from UCSC Genome Browser (http://www.genome.ucsc.edu/). Chromatin interactions were downloaded from 4Dgenome database (http://4dgenome.int-med.uiowa.edu). Transcription factor binding information were downloaded from ReMap database (http://pedagogix-tagc.univ-mrs.fr/remap). Dataset used in the study were summarised in (Extended Data Table S4).

### Statistical methods

Clinical phenotypes were summarized as mean (range: lowest-highest) or mean (percentage). All statistics for clinical phenotype was done with t-test (two-sided) or Wilcoxon Rank Test if t-test was not applicable. Detailed statistical methods were briefly denoted in the figure legends or text accompanying. All statistical analysis in this study were performed using R (3.5.1, 3.6.1 or 3.6.2) (http://CRAN.R-project.org).

### Molecular Biology

#### Nucleic acid preparation

Placenta genomic DNA was extracted from placenta tissue with Qiagen Animal Tissue DNA Extraction Kit (Qiagen 69504) according to the manufacturer's protocol. Cell-free DNA from serum was prepared using MagMAX Cell-Free DNA Extraction Kit (Thermo Fisher Scientific). Genomic DNA from FFPE tissue slides were extracted using MagPure Tissue DNA DF Kit (Magen Inc., MD5112-TL-06). Extracted DNA were quality-controlled by Qubit dsDNA HS assay (Thermo Fisher Scientific) and Agilent 2100 Fragment Analyzer.

#### Chromatin accessibility sequencing (ATAC-seq) on placenta

20mg villous samples were minced using a double-sized douncer (Sigma, D8938) in 1xHB (0.25M sucrose, 0.06M KCl, 0.015M NaCl, 0.005M MgCl2, 0.01M Tris-HCl pH 7.5), added to 5ml trypsin and 40ul 5U/ul DNase I (Sigma, D5025) and digested in 37 degree Celsius for 45 mins, with two times of rotation in between to mix the reaction. The digested cells were then neutralized with equal volume DMEM (Thermo Fisher, 11995065) +10% FBS (Gibco, 16000044) and filtered through a 70um cell filter (BD Falcon, 352350). The homogenize was centrifuged at 500x g, 4 degree Celsius for 5 mins. The sedimented cells were then resuspended in 400ul 1xHB and washed once, transferred to 2ml LoBind Tube (Eppendorf) and washed again. Cells were counted using Trypan blue (Solarbio, Beijing, China). After quantification, the cells were then added to a 30%-40%-50% iodixanol (Sigma, D1556) gradient and centrifuged at 3000x g, 20 mins at 4 degree Celsius. The cell layer at 30%-40% interface was collected for library preparation. DNA library were prepared ('tagmentation') with a Tn5 transposase kit (Vazyme, TD501) using 1 million cells per reaction according to manufacturer's protocol. After tagmentation and PCR amplification, the sequencing library were quality-controlled with SYBR-green based qPCR using primers for house-keeping gene (GAPDH) promoter and gene desert (chr5: 105187030-105190000) before sequencing.

#### Single stranded DNA methylation sequencing library preparation

Placenta genomic DNA (200ng) or cell-free DNA (1-10ng) were bisulfite converted using EZ-DNA Methylation-Gold Kit (Zymo Research, D5006) according to the manufacturer's protocol. After conversion, the DNA were subjected to a single-stranded library preparation protocol *Tequila 7N* (Euler Technology). In brief, the DNA were end-repaired using Klenow fragment (NEB) and tailed with poly-A homopolymer using terminal deoxynucleotide transferase (Takara), ligated to a poly-T overhang adaptor using T4 DNA ligase (Enzymatics), and linearly amplified for 12 cycles using PhusionU (Thermo Fisher Scientific). The amplified linear products were then annealed to a 5' adaptor with 7bp 3' random nucleotide overhang and PCR-amplified using adaptor oligos (Sangon, Shanghai, China) and Phusion (Thermo Fisher Scientific), resulting in a library with proper Illumina sequencing adaptor ends ready for NGS.

#### Preeclampsia methylation NGS capture panel

We designed a probe set covering both the Differential Methylated Region (DMR) between placenta and nongravida female cell-free DNA, and the DMR between normal and preeclampsia placenta. Probes were 120bp-long DNA fragments with added 5' and 3' PCR primers. The probe oligos were synthesized (Twist Inc. CA) and PCR-amplified using Phusion and biotinylated oligos (Sangon, Shanghai, China). Sequencing was performed on the oligo library using Kapa Hyper-prep sequencing kit (KAPA Biosciences) to ensure evenness of coverage of all oligos in the pool. To validate the performance of NGS capture panel, standard specimen was prepared by mixing placenta genomic DNA and a mixture of nongravida female cell-free DNA into a gradient of fetal fraction from 0% to 100%. The capture panel reliably identify placenta-specific DNA methylation pattern (methylation haplotype) and could deduce fetal fraction above 1% when sequenced to 10 million paired-end reads.

##### Methylation-specific capture sequencing

Capture sequencing library was prepared from single-stranded bisulfite-converted DNA library. Hybridization was done with SeqCap EpiGiant Enrichment Probe (Roche, 07138911001) or the in-house preeclampsia methylation NGS capture panel oligos and SeqCap wash and binding buffers (Roche) following the manufacturer's protocol. After hybridization, the library was amplified using Phusion for 8 cycles and sequenced on Novaseq sequencer (Illumina, CA) to a target of 100M paired-end 150bp reads (SeqCap EpiGiant) or 10M paired-end 150bp reads (In-house preeclampsia methylation NGS capture panel).

#### Data analysis

##### ATAC, Cut-And-Run and ChIP sequencing data preprocessing

Raw paired-end open chromatin tagmentation (ATAC), Cut-And-Run and chromatin immunopercipitation (ChIP) sequencing data, either downloaded from NCBI SRA or directly from in-house sequencing, were mapped to human reference genome GRCh37+decoy using Bowtie2 (-k 10 --very-sensitive -X 2000) (https://github.com/BenLangmead/bowtie2). All unmapped reads, non-uniquely mapped reads, reads with low mapping quality (MAPQ < 20) and PCR duplicates were removed. For single-cell and embryonic ATAC-seq sequencing libraries, data were used as is. For in-house prepared ATAC-seq data, the data were quality-controlled by assessing insertion size (using an in-house R script) and TSS-enrichment (using an in-house R script with GenomicRanges package (https://github.com/Bioconductor/GenomicRanges) measuring the depth ratio at the promoter region (hg19 refFlat annotation from UCSC Genome Browser) (0bp of TSS vs. 1kbp +- of TSS). A QC-passed ATAC-seq library must have TSS enrichment of 6, mapped deduplicated sequencing fragments >= 20M PE reads, PCB1>0.9, PCB2>3 (https://www.encodeproject.org/pipelines). Enrichment peaks were determined by intersecting peaks found from MACS2 callpeak (-f BAMPE) (https://github.com/taoliu/MACS) and Genrich (-r -m 1 -j; for ATAC only; and standard parameter for ChIP and Cut-and-Run) (https://github.com/jsh58/Genrich). Quality control of ATAC-seq libraries including read length, V-plot and TSS-enrichment were done with custom R script and deeptools (https://github.com/deeptools/deepTools).

##### ATAC-seq IDR peak identification

To identify a minimum set of open chromatin region to compare between different developmental stages, we first identify reliable open chromatin regions on each stage, by identifying repeatable ATAC enrichment peaks between technical and biological repeats with IDR (https://www.encodeproject.org/software/idr). Reliable ATAC peaks from different set of data were converged with 1bp minimum overlap and extended to the largest width of overlapping peaks. Joining these operation results in a set of non-overlapping, varied-width peaks across the genome encompassing all reliable open chromatin region found in embryonic and placenta sequencing data.

#### Correlation of ATAC-seq signal

Coverage (RPKM) of ATAC-seq on IDR peaks were calculated for each sample and Pearson correlation coefficient were calculated for each pair of samples in R (3.6.2).

#### Measurement of difference of ATAC-seq peaks

Read coverage of sequencing library were collected over the repeatable ATAC enrichment peak mentioned above with Sambamba. Differential enrichment was performed with DESeq2 (https://github.com/mikelove/DESeq2) using standard parameters. For samples with few replicates, we adapted a general linear model approach for estimating difference following the method mentioned in Reilly et al 2015[10]. Differential ATAC peaks between preeclampsia and normal placenta were summarized in Extended Data Table S2.

#### Histone modification analysis

Histone modification peaks were identified by MACS2 using following parameters: H3K4me3/H3K4me1/H3K27ac: -g hs -nomodel -nolambda; H3K27me3: -g hs --broad --broad-cutoff 0.05 -nomodel -nolambda. Weak (summed RPKM <20) or irreproducible peaks were removed for further analysis. Histone modification peaks overlapping known ATAC enrichment peak were used for further analysis. Read coverage of histone modification sequencing library were collected over the repeatable ATAC-and-histone-modification enrichment peak mentioned above with Sambamba, and normalized to RPKM (reads per kilo million for library), and Z-normalized.

#### Motif finding and transcription factor footprint analysis

Transcription factor binding motifs were collected from JASPAR (vertebrate non-redundant core motifs) and TFBS. Motifs were extracted using HOMER package (http://homer.ucsd.edu/homer/). Raw ATAC-seq data were pre-processed using RGT-Hint (www.regulatory-genomics.org/hint) package with standard parameter (rgt-hint footprinting --atac-seq --paired-end) and matched to motifs (rgt-motifanalysis matching --organism=hg19). Differential transcription factor footprint was analyzed by Hint[11] using (rgt-hint differential --organism=hg19 --bc) on three independent pairs of biological replicates between preeclampsia and non-preeclampsia placenta.

#### Genomic element annotation

Genomic element annotation was done using ANNOVAR (http://annovar.openbioinformatics.org/ ) and bedtools, where applicable. Differences on any class of genomic element were computed using fisher's exact test.

#### Genomic region liftover

For genomic regions liftover between UCSC hg18, hg19, pantro2, mm9 and GRCh37, the liftover executable from UCSC Kent Utility (https://genome.ucsc.edu/cgi-bin/hgLiftOver) and the lift-over synteny chain files from UCSC Genome Browser were used. Target genome was always UCSC hg19.

#### DNA methylation sequencing data preprocessing

Raw bisulfite-converted DNA methylation sequencing data, either downloaded from NCBI SRA or directly from in-house sequencing, were processed using fastp (--trim-front2 20 -w 20) (https://github.com/OpenGene/fastp) and mapped to GRCh37+decoy reference genome using BWA-Meth (https://github.com/brentp/bwa-meth) using standard parameters. Mapped data were deduplicated and sorted using Sambamba (https://github.com/biod/sambamba) and Samblaster (https://github.com/GregoryFaust/samblaster). CpG-methylation level were extracted using Pile-O-Meth (https://github.com/dpryan79/MethylDackel) toolkit. For all libraries, conversion rate were quality controlled by CHH methylation level >99%. Basic statistics of in-house sequencing library were further quality-controlled by on-target rate and on-target coverage with bedtools (https://github.com/arq5x/bedtools), and duplication rate and mapping rate with Sambamba.

For mouse data[12] of extraembryonic tissue (ExE) methylation, the sequencing reads were similarly preprocessed and mapped to mm9 reference. ExE-specific *de novo* methylation region from Smith et al[12] were directly used. To compare methylation level on human homologous regions, these mouse methylation regions were lifted-over from mm9 to GRCh37.

For Chimpanzee data of sperm methylation, the processed CpG methylation level from Molaro et al[8] was directly used, with lift-over from hg18 and pantro2 to hg19.

#### Differential methylation analysis

CpG methylation level (beta: defined as reads of C nucleotide over total read coverage on a single C or G base on CpG loci) was measured for each CpG loci across the genome as mentioned above using Pile-O-Meth. For each loci, beta from preeclampsia or non-preeclampsia (including normal, gestational hypertension, and gestational diabetes) pregnant female were summarized in R (3.6.2) using an in-house script. Differentially methylated loci (DML) were defined as: 1) $P<0.01$ for t-test between preeclampsia- and non-preeclampsia individuals , 2) beta difference between preeclampsia and non-preeclampsia individuals > 0.1. For DML used in cell-free DNA analysis, we further require 3) $P<0.01$ for t-test between preeclampsia placenta and nongravida female cfDNA, or $P<0.01$ for t-test between non-preeclampsia placenta and nongravida female cfDNA such that placental and maternal origin of cell-free DNA could be identified through methylation level or pattern.

#### Differentially methylated region (DMR) analysis

Initial DMR candidate were made by merging within-100bp-apart DML. The average beta of each initial DMR were calculated as mean beta of all CpG encompassed in the DMR. This average beta was subjected to t-test and P<0.01 regions were selected as candidate 'seed' DMR. Segments of methylation difference level were computed using a circular binary segmentation approach on beta difference between preeclampsia and non-preeclampsia placenta with DNAcopy (https://github.com/veseshan/DNAcopy). K-means clustering was performed using R (3.6.2) on the methylation beta difference on each segment, and clusters of segments fully encompassed candidate 'seed' DMR were selected as true DMR candidate (Extended Data Table S3).

#### DNA methylation haplotype analysis

Without considering technical duplication, the methylation pattern carried on single DNA sequencing read represents a single combination of CpG loci methylation from a single cell. We consider such single-read-carried combination of methylation pattern as a 'methylation haplotype' from a single cell. A difference on a single CpG loci (DML) or a region (DMR) is an aggregate statistics of all sequenced reads from a mixture of cells. However, a methylation haplotype always is from a single cell. Because of this, we consider methylation haplotype as a preferred subject to analysis, especially for biosamples containing a low frequency of DNA from the cell-of-interest.

Bisulfite sequencing reads were processed using Pysam (https://github.com/pysam-developers/pysam) into CpG methylation patterns labeled as 0 (unknown), 1 (methylated, C base) or -1 (unmethylated, T base) on each CpG loci. All reads covering a DMR were combined into a matrix and processed by Numpy (https://numpy.org/). Unique combinations of CpG methylation patterns were extracted, and for each unique combination (methylation haplotype), the relative frequency of this haplotype was computed for preeclampsia and non-preeclampsia placenta. The likelihood of haplotype (relative frequency) from the two sets of samples were compared using Fisher's exact test. Haplotypes with P<0.01 in Fisher's exact test comparison and frequency ratio > 2 (preeclampsia over non-preeclampsia, or vice versa) were denoted as specific methylation haplotype.

We termed the combination of all-methylated or all-unmethylated CpG as 'methylated' or 'unmethylated' haplotypes, and other haplotypes as 'mixed' haplotypes'. We calculated the distribution of haplotype on each DMR, using a Gaussian Mixture Model with Expectation-Maximization approach in mixtools (https://cran.r-project.org/web/packages/mixtools) (Extended Data Figure 4). Most DMR were dominanted by a single class of haplotype. Hence, we separate the DMR into hypomethylated (dominated by unmethylated haplotype in preeclampsia placenta) or hypermethylated (dominanted by methylated haplotype in preeclampsia placenta). On most DMR, these haplotypes encompass >2 CpG loci.

#### Hierachical clustering and correlation of DMR to epigenetic profiles

Mean beta value of DMR were calculated as the mean beta of all DMR-encompassed CpG. Hierachical clustering of samples and DMR were done by *hclust* function in R (3.6.2) with Ward method. To compare histone modification or transcription factor binding, *GenomicRanges* package in R (3.6.2) was used.

#### Functional enrichment of genomic loci around DMR

DMR loci were passed to GREAT (http://great.stanford.edu/public/html/) to analyze for any functional enrichment defined by BH-adjusted hypergeometrical test P<0.05 and BH-adjusted binomial test P<0.05.

#### Single cell RNAseq Data Analysis

Single cell RNAseq data and meta data were collected from NCBI SRA. Data mapping and single cell clustering was done with Monocle3 (https://github.com/cole-trapnell-lab/monocle-release) with standard practises. Trophoblasts (according to the metadata) from the dataset were subjected to further analysis. Pseudotime analysis and expression profiling was done with Monocle2 using DMR-associated genes or ATAC-seq differential peak associated genes.
DMR-associated gene were defined as genes with a promoter or enhancer overlapping at least 1bp with a DMR, or in adjacent (50kbps) of a DMR. We defined groups of informative DMR from clustered, aligned principle components using mean beta value of DMR.
Enhancer of a particular gene is defined with existence of IM-PET linkage between gene promoter and the enhancer element, using Hi-C, IM-PET and ChIA-PET data from 4Dgenome (http://4dgenome.int-med.uiowa.edu).
The ATAC-seq differential peak associated genes were defined as genes with a promoter or enhancer overlapping with at least 1bp with a differential peak in ATAC-seq.
Expression pseudotime plot was done with Monocle2[14] using the genes associated with DMR and/or ATAC-seq in single-cell RNAseq data. Note we preferred Monocle2 to Monocle3 in this plot merely because of the easiness of plotting and there should be no substantial difference in data processing.

#### Pseudotime Analysis of Methylation Data

Single cell or bulk methylation sequencing (bisulfite sequencing: WGBS or RRBS) data and metadata were collected from NCBI SRA. In-house data were described as mentioned before. Data mapping was done with Monocle3 (https://github.com/cole-trapnell-lab/monocle3) with standard practises, with the mean beta value as ‘expression value’. Dispersion was estimated with negative binomial model. Data were preprocessed using 10 dimensions with UMAP. To cluster samples from different techniques, we used ‘align_cds’ function from Monocle3 using mutual nearest neighbor alignment method[13], and performed pseudotime trajectory analysis on these aligned data.

#### Pseudotime Analysis of ATAC-seq Data

Single cell or bulk ATAC sequencing data and metadata were collected from NCBI SRA. In-house data were described as mentioned before. The overall processing was similar to pseudotime analysis of methylation data mentioned above, with the only difference that ATAC-seq RPKM from all DMR regions were used as ‘expression level’ in the analysis.

#### Predicting placenta methylation using cell-free DNA methylation pattern

Cell-free DNA and placenta were subjected to methylation capture sequencing using the preeclampsia methylation NGS capture panel mentioned above. Data was processed and subjected to methylation haplotype extraction using Numpy. After extraction, a table of methylation haplotype frequency was formed for each sample. For the placenta model, haplotype frequency from preeclampsia and non-preeclampsia placenta sequencing data were subjected to principle component analysis (PCA) using R (3.6.2). The resulted PCA predicted component weights were modelled with general linear model (GLM) using glmnet (https://cran.r-project.org/web/packages/glmnet/index.html) with target index value 0=preeclampsia and 100=non-preeclampsia. Such prediction results in a score we termed "developmental maturity" which could be expressed in arbitrary units. The cell-free DNA data from preeclampsia and non-preeclampsia pregnant female were pre-processed with the trained PCA model and further predicted using the GLM. The result GLM predictive value indicates non-preeclampsia placenta methylation pattern load from cell-free DNA.

#### Predicting pregnancy outcome from cell-free DNA methylation data

A general linear model was derived to predict risk of preeclampsia using the same PCA as described above and target index value 1=preeclampsia and 0=non-preeclampsia. AUC (https://cran.r-project.org/web/packages/AUC) analysis was performed to find optimal cut-off threshold for GLM model predicting preeclampsia v.s. non-preeclampsia cell-free DNA data. To validate the model, cell-free DNA from preeclampsia and non-preeclampsia pregnant female in the clinical validation cohort were subjected to methylation capture sequencing using the preeclampsia methylation NGS capture panel mentioned above, and processed using a similar model. Clinical information was unblinded after prediction was made. Predictive sensitivity and specificity (kappa) was done by standard practises.

### Supplementary Reference

1. Wisner, K. Gestational Hypertension and Preeclampsia. *MCN Am. J. Matern. Nurs.* **44**, 170 (2019).
2. Liu, L. *et al.* An integrated chromatin accessibility and transcriptome landscape of human pre-implantation embryos. *Nat. Commun.* **10**, 1–11 (2019).
3. Wu, J. *et al.* Chromatin analysis in human early development reveals epigenetic transition during ZGA. *Nature* **557**, 256–260 (2018).
4. Zhu, P. *et al.* Single-cell DNA methylome sequencing of human preimplantation embryos. *Nat. Genet.* **50**, 12–19 (2018).
5. Xia, W. *et al.* Resetting histone modifications during human parental-to-zygotic transition. *Science (80-. ).* **365**, 353–360 (2019).
6. Vento-Tormo, R. *et al.* Single-cell reconstruction of the early maternal–fetal interface in humans. *Nature* **563**, 347–353 (2018).
7. Gamage, T. K. J. B. *et al.* Human trophoblasts are primarily distinguished from somatic cells by differences in the pattern rather than the degree of global CpG methylation. *Biol. Open* **7**, (2018).
8. Molaro, A. *et al.* Sperm methylation profiles reveal features of epigenetic inheritance and evolution in primates. *Cell* **146**, 1029–1041 (2011).
9. Gao, L. *et al.* Chromatin Accessibility Landscape in Human Early Embryos and Its Association with Evolution. *Cell* **173**, 248-259.e15 (2018).
10. Reilly, S. & Yin, J. Evolutionary Changes in Promoter and Enhancer Activity During Human Corticogenesis Steven. *Science (80-. ).* **347**, 1155–1159 (2015).
11. Li, Z. & Schulz, M. Identification of transcription factor binding sites using Gaussian mixture models. *Genome Biol.* **31**, 70–80 (2014).
12. Smith, Z. D. *et al.* Epigenetic restriction of extraembryonic lineages mirrors the somatic transition to cancer. *Nature* **549**, 543–547 (2017).
13. Haghverdi, L., Lun, A. T. L., Morgan, M. D. & Marioni, J. C. Batch effects in single-cell RNA-sequencing data are corrected by matching mutual nearest neighbors. *Nat. Biotechnol.* **36**, 421–427 (2018).
14. Qiu, X. *et al.* Reversed graph embedding resolves complex single-cell trajectories. *Nat. Methods* **14**, 979–982 (2017).

### Legends for Extended Data Figure

**Extended Data Figure 1. Robustness and validity of ATAC-seq from placenta**
**a**: Placenta ATAC-seq signals overlap with activating histone mark (H3K27ac). Bulk placenta ATAC-seq data in this study shown as dark grey and H3K27ac ChIP-seq data of placenta from Roadmap Project (GSM1127147) shown as red. Majority of H3K27ac peaks overlapped with placenta ATAC-seq peaks, shown in light orange shades, indicating that placenta ATAC-seq peaks marked open chromatin with active transcription.
**b**: Open chromatin accessibility distinguishes early developmental stages of human embryos and pathological state of placentas. Per-sample correlation of single cell ATAC-seq peaks from zygote[2,3], 2-cell[3], 4-cell[2,3] 8-cell[2,3], morula[2], ICM[3], trophectoderm (TE) [2] hESC[2,3], and bulk placenta ATAC-seq peaks from preeclampsia and non-preeclampsia placenta were shown. Overall, samples of similar developmental stage and pathological state were clustered together. Samples from pre-ZGA, and post-ZGA, clustered together regardless of study origin, suggesting that these ATAC-seq data were sufficiently robust and batch effect were successfully alleviated. Preeclampsia and non-preeclampsia placenta clustered separately. Preeclampsia placentas (placenta PE, black) had distinctive ATAC peak profile comparing with non-preeclampsia placentas (placenta and placentaProU). Placentas from pregnancy with proteinuria but not preeclampsia (placentaProU, grey) failed to distinguish with placentas from normal pregnancy (placenta, white) based on bulk ATAC-seq peaks. Labels on the side: tagtype: embryonic developmental stage or pathological state. ICM: inner cell mass, TE: trophectoderm, differentiated_cells: cells differentiated from hESC; Placenta: non-preeclampsia normal placenta; PlacentaProtU: placenta from proteinuria, non-preeclampsia pregnancy; PlacentaPE: placenta from preeclampsia pregnancy. PN: pronucleus (for embryonic development data, as in the original publication).
**c**: V-plot around CTCF binding site on an example placenta ATAC-seq data, showing characteristic histone and CTCF binding footprint.
**d**: Agilent Bioanalyzer 2100 run of an example placenta ATAC-seq library, showing characteristic peaks of fragment length at ~200bp (open chromatin), ~380bp (mononucleosome), ~550bp (dinucleosome).
**e**: Read length distribution from an example placenta ATAC-seq data, showing characteristic peaks of fragment length at open chromatin, mononucleosome, dinucleosome, and trinucleosome.
**f**: TSS-enrichment (sequencing depth around +- 2kbp of transcription start site) of an example placenta ATAC-seq library, showing enrichment at 0bp of TSS >30-fold of boundary (+-2kbp).

**Extended Data Figure 2**. **Chromatin accessibility distinguishes preeclampsia to normal placenta**

(a-f) Preeclampsia placentas (PE) exhibit differential ATAC profiles at *PAPPA2*, *FLT1*, *LIFR*, *KDR*, *VEGFA* and *PGF*, comparing with placentas from non-preeclampsia pregnancies (CTRL). Light blue shades represent ATAC peaks that are lessened or lost in PE and light green shades show ATAC peaks that are increased in PE. Red lines represent preeclampsia-DMR regions. Combined ATAC-seq signals from 2-cell, 4-cell, 8-cell, hESC were also shown in the figures for comparison. (g/h) Enrichment of significantly ATAC-seq peaks on Reactome Pathways from PantherDB. Significantly enriched pathway with FDR < 0.01 were shown on the figure. X axis: log P value (unadjusted). (g) pathways associated with preeclampsia-gained peaks with significant differences; (h): pathways associated with preeclampsia-lost peaks.

**Extended Data Figure 3**. **Differential methylated regions between preeclampsia and normal placenta across the genome.**
Preeclampsia-hyper-methylated (red) and hypo-methylated (blue) regions were shown on chromosome.

**Extended Data Figure 4. Preeclampsia-specific DMR were dominanted by either hypermethylated or hypomethylated haplotypes.**
a: Most DMR were dominanted by either hypermethylated or hypomethylated haplotype. x axis: fraction of 'fully methylated' state-associated (normal or preeclampsia-specific) methylation haplotype on each DMR window; y axis: DMR window number.
b: Distribution of hypermethylated (meth fraction > 0.5, blue) or hypomethylated (meth fraction <0.5, red) DMR on each chromosome.
c: Distribution of CpG number in each DMR for hypomethylated (blue), hypermethylated (red) and mixed (white) DMR. y axis: CpG number contained by state-specific methylation haplotype in the DMR.

**Extended Data Figure 5. Mean methylation level of preeclampsia specific DMR could distinguish preeclampsia placenta to non-preeclampsia placenta**
x axis: Mean beta of preeclampsia-hypermethylated DMR from each sample; y axis: Mean beta of preeclampsia-hypomethylated DMR from each sample. Single cell WGBS from oocyte, sperm, 8-cell stage, morula, ICM, 6-week fetus, and bulk genomic DNA MCBS of placenta from normal (CTRL), twin-twin transfusion-syndrome (TTTS), gestational diabetes (GDM), gestational hypertension (GHT), or preeclampsia pregnancy (PE) were shown in the figure. Mean beta were calculated as the mean methylated CpG fraction in DMR. As shown in the figure, most PE placenta could be distinguished from all other placentas by hypermethylated PE-hyper DMR and hypomethylated PE-hypo DMR.

**Extended Data Figure 6. Enrichment of TFBS in DMR groups.**
DMR fine groups (shown on the right of panel) were classified as in Figure 1. Peaks from transcription factor chromatin immunoprecipitation sequencing (ChIP-seq) data were collected (from ReMap 2018) and overlapped with DMR. Different DMR fine groups showed differential enrichment of TFBS. Polycomb binding (EZH2, SUZ12, EED, JARID2, YY1, CBX1) were found in both PE-hypo (group 1/10/11) and PE-hyper (group 5/6/8) DMRs, suggesting that lineage specification and cell fate determination might be changed in preeclampsia. CTCF binding sites were enriched in PE-hypo DMR (group 1/11), suggesting that topological associated domain might be changed in preeclampsia because CTCF binding to DNA is dependent on CpG de-methylation of core CTCF binding motif. HIF1A and immediate-early-gene (IEG: FOS/JUN/JUND) binding sites were enriched in PE-hypo DMR, suggesting that PE-hypo DMR might dynamically respond to hypoxia and cellular stress.

**Extended Data Figure 7. Pathway enrichment of DMR groups.**
GREAT (http://great.stanford.edu/public/html/) enrichment of DMR loci were shown in the figure. PcG-enriched DMR groups (Group 6 /G6 and Group 11 /G11) were functionally associated with pathways involved in pattern specification (both), embryo development (G6), regionalization (G6), and cell fate commitement (G11), suggesting that these loci were involved in control of lineage specification genes.

**Extended Data Figure 8. Preeclampsia placenta failed to develop the cancer-like somatic *de novo* methylation as in normal placenta.**

a/b) Box-and-whisker plot of methylation level (beta) of mammalian conserved PE-hypo DML on individual samples; light blue: normal samples; dark blue: tumor samples. light green: preeclampsia fetal surface placenta samples; dark green: normal fetal surface placenta samples. c/d) Box-and-whisker plot of methylation level (beta) of non-conserved, human-specific PE-hypo DML on individual samples; light blue: normal samples; dark blue: tumor samples. light green: preeclampsia fetal surface placenta samples; dark green: normal fetal surface placenta samples. e/f) Box-and-whisker plot of methylation level (beta) of all PE-hyper DML on individual samples; light blue: normal samples; dark blue: tumor samples. light green: preeclampsia fetal surface placenta samples; dark green: normal fetal surface placenta samples.

**Extended Data Figure 9. Differential functional element enrichment of preeclampsia-associated DMR**

a) Preeclampsia-hypermethylated (PE-hyper) and preeclampsia-hypomethylated (PE-hypo) DMR were enriched differentially with genomic functional elements. Comparing to PE-hyper sites, PE-hypo sites were found more in 5'UTR, intronic and exonic regions, suggesting regulatory function of PE-hypo DMR. b) Mammalian-conserved ExE-de-novo methlyation loci and human-specific ExE-de-novo methylation loci in TSS were hypomethylated in preeclampsia placenta. c) LTR12C with sperm-specific hypomethylation (sperm HMR) in human were hypermethylated in preeclampsia placenta. y axis: per-CpG methylation difference (PE-CTRL). d/e) Example sperm HMR (red: common, green: human-specific, not sperm HMR in chimpanzee) encompassing preeclampsia-hypermethylated LTR12C around *GCNT2* and *APOBEC1*. Black bars above: Methylation level (beta) of CpG loci in chimpanzee (top) or human (bottom) genomes. Chimpanzee data were lifted to human genome by LiftOver. Sperm-specific hypomethylation valley could be immediately identified visually on the figure. LTR12C loci were shown next to genes. Enlarged panels showing per-CpG methylation difference (PE-CTRL) of each LTR12C loci in the sperm hypomethylation valley of human.

**Extended Data Figure 10. Mean methylation level of LTR12C-carried DMR could distinguish preeclampsia placenta to non-preeclampsia placenta**

x axis: Classes of different retrotransposons of DMR; y axis: Mean beta of retrotransposon (different class) in each sample. a: Methylation level of LTR12-family ERV (LTR12C, LTR12D, HERV9-int) distinguishes preeclampsia fetal side placenta (light blue) from normal fetal side placenta (dark blue). b: Methylation level of primate-specific ERV (labelled as chimp and human icon) differentiates preeclampsia and normal placenta.

**Extended Data Figure 11. Single cell RNAseq pseudotime trajectory of trophoblast**

Trophoblast single cell RNAseq result were subjected to dimentionality reduction in Monocle3 using UMAP and projected into low dimention space. Pseudotime trajectory from villous cytotrophoblast (VCT) towards syncytotrophoblast (SCT) or extravillous trophoblast (EVT) were inferred using Monocle3. Cells were colored according to phenotype (EVT: gold; VCT: purple; SCT: blue).

**Extended Data Figure 12. LTR12C hypermethylation in preeclampsia placenta**

Preeclampsia-hypermethylated LTR12C around *PJA2*, *IER2*, *SLC30A8*, *GABARAPL1*, *GCNT2* and *PAPPA2* were shown in the figure. Top: ATAC-seq of 2-cell, 4-cell, 8-cell, ICM, hESC, PE and CTRL Placenta. Bottom: Methylation difference (beta: PE-CTRL) on the loci, shown together with LTR12C genomic direction. These data showed that preeclampsia-hypermethylated LTR12C were commonly hypermethylated in 5' transcriptional start regions. These LTR12C were most active around ZGA (8-cell stage) and silenced post-ZGA.

**Extended Data Figure 13. Dynamics of preeclampsia-hyper-DMR (PE-hyper-DMR) and preeclampsia-hypo-DMR (PE-hypo-DMR) methylation level across different stages of embryonic development**

a: Methylation profile of PE-hyper- and PE-hypo- DMR in oocyte. DNA methylation on both PE-hyper-DMR and PE-hypo-DMR loci exhibit bimodal pattern in oocyte.

b: Methylation profile of PE-hyper- and PE-hypo- DMR in sperm. PE-hyper-DMR loci are hypo-methylated in sperm. PE-hypo-DMR loci exhibit bimodal methylation pattern in sperm.

c: Methylation profile of PE-hyper- and PE-hypo- DMR at the time of fertilization (predicted by oocyte+sperm/2).

d: Methylation profile of PE-hyper- and PE-hypo- DMR at 8-cell stage. PE-hyper-DMR loci are hypomethylated at 8-cell stage while PE-hypo-DMR loci show bimodal methylation pattern.

e: Methylation profile of PE-hyper- and PE-hypo- DMR at morula stage. PE-hyper-DMR loci are hypomethylated at morula stage while PE-hypo-DMR loci show bimodal methylation pattern.

f: Methylation profile of PE-hyper- and PE-hypo- DMR in ICM. In ICM, partially methylation on PE-hyper- and PE-hypo- DMR is increased.

g: Methylation profile of PE-hyper- and PE-hypo- DMR in non-preeclampsia placenta (CTRL placenta). In CTRL placenta, both PE-hyper- and PE-hypo- DMR are hypermethylated.

h: Methylation profile of PE-hyper- and PE-hypo- DMR in preeclampsia placenta (PE placenta). Comparing with CTRL placenta, PE placentas show increased methylation level at PE-hyper-DMR loci and decreased methylation level at PE-hypo-DMR loci.

i: The methylation profile of PE-hyper- and PE-hypo-DMR at sperm stage (y axis) comparing with oocyte stage (x axis) suggesting that PE-hypo loci were hypermethylated in both gamete but all PE-hyper loci were hypomethylated in sperm.

j: The methylation profile of PE-hyper- and PE-hypo-DMR at 8-cell stage (y axis) comparing with zygote stage (x axis) suggesting that both PE-hyper and PE-hypo loci undergone half demethylation at 8-cell stage.

k: The methylation profile of PE-hyper- and PE-hypo-DMR at morula stage (y axis) comparing with 8-cell stage (x axis).

l: The methylation profile of PE-hyper- and PE-hypo-DMR in ICM (y axis) comparing with morula stage (x axis).

m: The methylation profile of PE-hyper- and PE-hypo-DMR in CTRL placenta (y axis) comparing with ICM (x axis) suggesting that placenta exhibited de novo methylation on these loci.

n: The methylation profile of PE-hyper- and PE-hypo-DMR in PE placenta (y axis) comparing with ICM (x axis).

o: The methylation profile of PE-hyper- and PE-hypo-DMR in PE placenta (x axis) comparing with CTRL placenta (y axis).

**Extended Data Figure 14. LTR12C hypermethylation in preeclampsia placenta affects gene transcription and topological associated domain.**

Preeclampsia-hypermethylated LTR12C around *PAPPA2* (a), *CDKN3* (b) and *YY1* (c) were shown in the figure. Top: ATAC-seq of CTRL and PE Placenta. Middle: ChIP-seq of H3K9me3, H3K27me3, H3K27ac, H3K4me1 and H3K4me3 in placenta (Roadmap

Consortium) showing that the hypermethylated LTR12C could be associated with repressive histone marks such as H3K9me3 (a) or H3K27me3 (b) but not activating histone marks. Bottom: chromatin cis-interaction frequency and detail links between loci on the region (4Dgenome) showing that hypermethylated LTR12C were functinoally relevant within topological associated domains (TAD). For *PAPPA2* (a), which was up-regulated in preeclampsia, gain-of-expression of the gene is associated with gained chromatin accessibility as well as a hypermethylated LTR12C in its neighbouring TAD. For *YY1* and *CDKN3*, which were down-regulated in preeclampsia, downregulation of the gene is associated with less chromatin accessibility on their promoter and enhancer regions, and hypermethylated LTR12C linking to their enhancer/promoters in the same TAD. These results suggest that LTR12C methylation might switch the gene expression dynamics in its relevant TAD.

**Extended Data Figure 15. Preeclampsia-hypomethylated DMR**
Preeclampsia-hypomethylated DMR around *DOCK5*, *FGFR3*, *E2F8*, *LRRFIP1*, *LIMS2* and *TMP4* were shown in the figure. Top: ATAC-seq of 2-cell, 4-cell, 8-cell, ICM, hESC, PE and CTRL Placenta. Bottom: Methylation difference (beta: PE-CTRL) on the loci, shown together with LTR12C genomic direction. These data showed that preeclampsia-hypomethylated DMR were only active in post-ZGA stages.

**Extended Data Figure 16. Pseudotime evolution trajectory built with whole-genome methylation profiles on DMR across embryonic development stages**
Mean beta level from preeclampsia-hypomethylated DMR were used to build pseudotime evolution trajectory with single cell WGBS data of different embryonic developmental stages from oocyte, sperm, zygote, 2-cell stage, 4-cell stage, 8-cell stage, morula, trophectoderm (TE), inner cell mass (ICM), undistinguished blastocyst, as well as bulk RRBS data from extravillous trophoblast (EVT), spongiotrophoblast (SPT), cellular trophoblast (CTB), and bulk MCBS data from fetal-side (facetofetus) and maternal side (facetomother) of placenta of normal (CTRL) or preeclampsia (PE) pregnancy. The trajectory suggested delay of preeclampsia placenta during the development from trophectoderm to trophoblasts.

**Extended Data Figure 17. Chromatin accessibility on DMR across embryonic development stages**
Mean chromatin accessibility level from differential ATAC peak on preeclampsia-specific DMR were shown with single cell ATAC-seq data of different embryonic developmental stages from oocyte, sperm, zygote, 2-cell stage, 4-cell stage, 8-cell stage, blastocyst, ESC (ground state and primed state), and bulk ATAC-seq data from trophoblasts and placentas (CTRL or PE).

**Extended Data Figure 18. Preeclampsia-specific methylation haplotype in *FLT1*-associated preeclampsia-specific enhancer**
Preeclampsia-specific methylation haplotypes with lower methylation level (per-read CpG methylation frequency statistics for placenta and plasma cell-free DNA in inset bar graphs) on *FLT1* preeclampsia-specific enhancer detected in cell-free DNA.

**Extended Data Figure 19. Functional enrichment of DMR with chromatin accessibility differences**
(a) Enrichment of chromatin-accessibility-difference-overlapped-DMR on ChromHMM regions. (b) Enrichment of chromatin-accessibility-difference-overlapped-DMR associated genes on Reactome Pathways from PantherDB. Significantly enriched pathway with FDR <

0.01 were shown on the figure. X axis: log P value (unadjusted). Left: all loci; middle: upregulated (preeclampsia-gain) loci; right: downregulated (preeclampsia-loss) loci.

### Legends for Extended Data Table

Extended Table S1: Clinical information of placentas used in this study. * $p<0.05$ in chi-square test.
Extended Table S2: Table of differential ATAC peaks in PE and Normal
Extended Table S3: Table of DMR in PE and Normal
Extended Table S4: Data used in the study
Extended Table S5: Software version used in the study
Extended Table S6: Clinical information of retrospective validation cohort. * $p<0.05$ in chi-square test.
