## Supplementary Data 1b Placenta Donor Characteristics for "Developmentally Delayed Epigenetic Reprogramming Underlying the Pathogenesis of Preeclampsia"

|  | **Preeclampsia** | **non-Preeclampsia** |
| --- | --- | --- |
| **Age (years, range)** | 34.75 (30-41) | 32.33 (25-40) |
| **Body Mass Index (kg/m^2^, range)** | 24.88 (19.68-35.29) | 22.62 (17.04-30.49) |
| **Gestation (times, range)** | 2.92 (2-4) | 2.48 (1-5) |
| **Parity (times, range)** | 1.58 (1-2) | 1.81 (1-4) |
| **Assisted reproduction (%)** | 8.30 (1/12) | 8.70 (2/23) |
| **Labour week (weeks, range)** | 34.75 (29-38) | 38.22 (31-40) |
| **Previous adverse pregnancy outcome (%)** | 0.00 (0/12) | 8.70 (2/23) |
| **Chronic diseases (%)** | 41.67 (5/12) | 0.00 (0/23) |
| Hypertension | 16.67 (2/12) | 0.00 (0/23)* |
| Pregestational diabetes | 25.00 (3/12) | 0.00 (0/23)* |
| Autoimmune disease | 8.30 (1/12) | 0.00 (0/23) |
| G6PD deficiency | 8.30 (1/12) | 0.00 (0/23) |
| **Pregnancy complications (%)** | 100.00 (12/12) | 43.48 (10/23) |
| Intrahepatic cholestasis of pregnancy | 25.00 (3/12) | 0.00 (0/23)* |
| Gestational hypertension | 100.00 (12/12) | 21.74 (5/23)* |
| Gestational diabetes mellitus | 8.30 (1/12) | 26.09 (6/23) |
| Preterm birth | 41.67 (5/12) | 8.70 (2/23)* |
| Placenta adhesion | 8.30 (1/12) | 0.00 (0/23) |
| Battledore placenta | 8.30 (1/12) | 0.00 (0/23) |
| Velamentous placenta | 0.00 (0/12) | 0.00 (0/23) |
| Placenta restriction | 6.25 (1/16) | 0.00 (0/23) |
| Postpartum Haemorrhage | 8.30 (1/12) | 0.00 (0/23) |
| Fetal growth restriction | 16.67 (2/12) | 4.35 (1/23) |
| Huge baby | 0.00 (0/12) | 4.35 (1/23) |

Extended Data Table 1b. Clinical profile of pregnancies in analysing placentas
