## Supplementary Data 6 Retrospective Clinical Cohort Characteristics for "Developmentally Delayed Epigenetic Reprogramming Underlying the Pathogenesis of Preeclampsia"

|  | **Preeclampsia** | **non-Preeclampsia** |
| --- | --- | --- |
| **Age (years, range)** | 32.72 (22-41) | 32.35 (21-42) |
| **Body Mass Index (kg/m^2^, range)** | 23.03 (15.63-33.33) | 21.10 (16.00-31.64) |
| **Gestation (times, range)** | 2.36 (1-7) | 2.21 (1-6) |
| **Parity (times, range)** | 1.48 (1-2) | 1.63 (1-3) |
| **Assisted reproduction (%)** | 10.81 (4/37) | 2.46 (3/122) |
| **Labour week (weeks, range)** | 36.47 (28-40) | 38.93 (37-41) |
| **Adverse pregnant history (%)** | 13.51 (5/37) | 4.92 (6/122) |
| **Chronic diseases (%)** | 16.98 (9/53) | 1.64 (2/122) |
| Hypertension | 5.41 (2/37) | 0.00 (0/122)* |
| Pregestational diabetes | 0.00 (0/37) | 0.00 (0/122) |
| G6PD deficiency | 2.70 (1/37) | 0.00 (0/122) |
| Coagulation dysfunction | 5.41 (2/37) | 0.00 (0/122)* |
| Kidney disease | 8.11 (3/37) | 0.00 (0/122)* |
| Impaired liver function | 8.11 (3/37) | 0.00 (0/122)* |
| Autoimmune disease | 0.00 (0/37) | 1.64 (2/122) |
| **Pregnancy complications (%)** | 100.00 (37/37) | 32.79 (40/122) |
| Intrahepatic cholestasis of pregnancy | 2.70 (1/37) | 0.00 (0/122)* |
| Gestational hypertension | 100.00 (37/37) | 3.28 (4/122)* |
| Gestational diabetes mellitus | 29.73 (11/37) | 1.64 (2/122)* |
| Preterm birth | 24.32 (9/37) | 0.00 (0/122)* |
| Postpartum haemorrhage | 5.41 (2/37) | 0.00 (0/122)* |
| Placenta adhesion | 8.11 (3/37) | 3.28 (4/122) |
| Prelabour rupture of the membranes | 18.92 (7/37) | 12.30 (15/122) |
| Battledore placenta | 8.11 (3/37) | 1.64 (2/122) |
| Velamentous placenta | 0.00 (0/37) | 2.46 (3/122) |
| Placenta previa | 0.00 (0/37) | 0.82 (1/122) |
| Fetal growth restriction | 29.73 (11/37) | 12.30 (15/122)* |
| Small for gestational age | 8.11 (3/37) | 0.82 (1/122)* |
| Huge baby | 5.41 (2/37) | 4.92 (6/122) |

Extended Data Table 6. Clinical profile in retrospective study
